# Prediction of the 3D cancer genome from genomic rearrangements using InfoHiC

**DOI:** 10.1101/2022.08.02.502462

**Authors:** Yeonghun Lee, Sung-Hye Park, Hyunju Lee

## Abstract

Although cancer genomes often contain complex genomic rearrangements, its impact on tumorigenesis is still unclear, especially when they are involved in non-coding regions. Understanding 3D genome architecture is crucial for uncovering the impacts of genomic rearrangements. Here, we present InfoHiC, a method for predicting 3D genome folding and cancer Hi-C from complex genomic rearrangements. InfoHiC provides distinct interaction views of multiple contigs from the cancer Hi-C matrix. We then validated cancer Hi-C prediction using breast cancer cell line data and found contig-specific interaction changes. Moreover, we applied InfoHiC to patients with breast cancer and identified neo topologically associating domains and super-enhancer hijacking events associated with oncogenic overexpression and poor survival outcomes. Finally, we applied InfoHiC to pediatric patients with medulloblastoma, and found genomic rearrangements in non-coding regions that caused super-enhancer hijacking events of medulloblastoma driver genes (*GFI1, GFI1B*, and *PRDM6*). In summary, InfoHiC can predict genome folding changes in cancer genomes and may reveal therapeutic targets by uncovering the functional impacts of non-coding genomic rearrangements.

## Introduction

Complex genomic rearrangements have been observed in various cancers^1,2^. Moreover, high-throughput sequencing revealed breakpoints of genomic rearrangements or so-called structural variations (SVs) at the single-base level, and revealed an association with tumorigenesis in terms of copy number alterations (CNAs) or gene fusions^3^. A recent study revealed complex genomic rearrangements including homogeneously staining regions (HSRs), double minutes (DMs), and breakage-fusion-bridge cycles, which contributed to massive gene copy number (CN) changes^1^. However, most SV breakpoints have been found in non-coding regions or outside genes, and their effect on tumorigenesis has been less closely investigated.

Recent studies^4–6^ using Hi-C data have advanced our understanding of the 3D genome organization of DNA sequences. Topologically associating domains (TADs)^7^ are key components of the 3D genome architecture, and DNA elements in the same TAD are more likely to interact with each other. SVs may form neo-TADs and novel interactions between gene promoters and enhancers^6^. However, neo-TAD formation has been less thoroughly investigated because of a lack of available methods for interpreting cancer Hi-C data. Recently, a method called NeoLoopFinder^8^ was developed to analyze neo-loops from cancer Hi-C data; however, this method is restricted to large SVs, and interactions from complex SV clusters are ignored. In addition, high-quality Hi-C data are limitedly available in cancer cell lines; thus, the 3D genome architecture of primary cancers has not yet been investigated. Furthermore, although deep learning approaches^9–11^ have been used to predict Hi-C data using sequence-based modeling, these methods are based on the reference sequence and are restricted to non-cancerous cell lines. Their applicability in cancer Hi-C data were not investigated yet.

In this article, we present a method named InfoHiC that enables cancer Hi-C prediction from complex SVs. Based on complex genomic rearrangements in different haplotypes, InfoHiC predicts the cancer Hi-C matrix in assembled contig and total Hi-C views. We then validate our model using Hi-C data from breast cancer cell lines (T47D, BT474, HCC1954, SKBR3, and MCF7) and show that InfoHiC outperforms the reference-based model. We also discover neo-TADs and neoloops, as well as enhancer hijacking events from contig Hi-C prediction in breast cancer cell lines, which can be validated using the InfoHiC validation scheme. Furthermore, we apply InfoHiC to patients with breast cancer from The Cancer Genome Atlas (TCGA) and discover enhancer hijacking events that result in oncogenic overexpression and poor survival outcomes. Finally, we apply InfoHiC to pediatric patients with medulloblastoma, and highlight its analytical potential for discovering non-coding driver SVs associated with oncogenic expression.

## Results

### InfoHiC predicts cancer Hi-C using a convolutional neural network (CNN) model

A cancer Hi-C matrix is a mixture of Hi-C matrices from multiple genomic contigs, in which Hi-C reads from different contigs are observed as a sum in the reference coordinates. In contrast to reference-based models^9–11^, the cancer Hi-C prediction model requires merging of multiple prediction outputs. In addition, contigs have different genomic variations, including single nucleotide polymorphisms (SNPs), SVs, and CNAs; thus, genomic variations must be encoded in a contig matrix. To this end, we developed an architecture called InfoHiC, composed of CNNs, that outputs chromatin interactions of genomic contigs and merges them in the total Hi-C views based on haplotype-specific copy number (HSCN) encoding, which represents genomic variants in the contig matrix (Fig. 1).

**Fig. 1:**
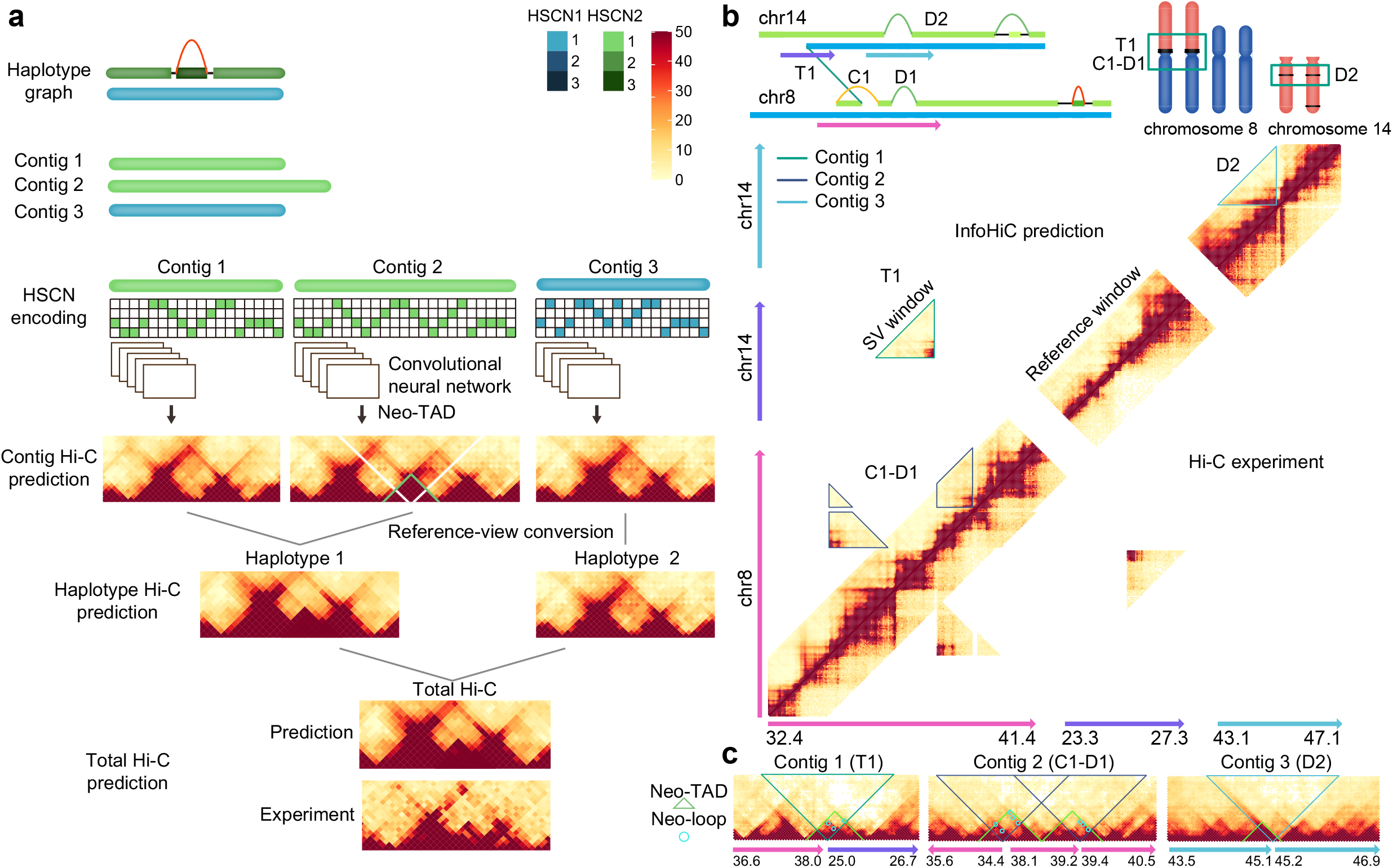
Schematic diagram of InfoHiC. **a**, InfoHiC prediction of the total Hi-C matrix from multiple genomic contigs. Genomic contigs were derived from the haplotype breakpoint graph (haplotype 1 in blue and haplotype 2 in green), and the HSCN matrix was employed to encode the SNP composition and CN of each contig. contig 2 has tandem duplication, and the CNN predicts a neo-TAD (green) from contig 2. Hi-C predictions from contig 1, 2, and 3 were successively merged into the haplotype Hi-C and the total Hi-C matrix. **b**, InfoHiC prediction of chromosomes 8 and 14 in the T47D cell line. SVs (T1, C1, D1, and D2) are annotated in the haplotype graph and karyotypes with two copies of der(8)t(8;14). The upper diagonal matrix represents the InfoHiC prediction and the lower diagonal matrix represents the Hi-C experiment of T47D. SV-induced Hi-C contacts are outlined by each genomic contig. **c**, Contig Hi-C prediction by InfoHiC. Three contig Hi-C matrices were annotated with SVs (T1, C1-D1, and D2), neo-TAD (green), and neoloop (cyan) annotation; reference coordinates are shown below at the mega-base scale.

To predict a target pair of the Hi-C contact between genomic bins observed in the reference coordinates, we collected contig DNA sequences in which a contig target pair of genomic bins (40 kb) is matched with the reference target pair in a pre-defined genomic window with a length *l* of 1 Mb or 2 Mb. For the contig, we used InfoGenomeR^1^, which enables the assembly of haplotype contigs from SVs and SNPs, and measures their CNs using whole-genome sequencing (WGS) data. Next, based on haplotype contigs, we encoded contig DNA features using an 4 *× l* matrix, where the row index represents the HSCN of each base sequence (A, C, G, and T). We used a CNN model as a component of the sequence-based prediction for each genomic contig, which has a similar architecture with deepC^9^ that is developed for a single reference sequence. Each CNN model predicts a target contact, and target contacts may lie in different distances in the contig coordinates. The predicted contig Hi-C matrices are then transformed into reference coordinate matrices and merged into the haplotype Hi-C and the total Hi-C matrix (Fig. 1a).

Moreover, we propose a validation scheme of 3D genome folding changes predicted from SVs using InfoHiC, which maps contig Hi-C matrices into the reference coordinates and validates them via a cancer Hi-C experiment (Fig. 1b). Scattered Hi-C contacts resulting from SVs are outlined by each genomic contig and those from the InfoHiC prediction and the Hi-C experiment are compared in the reference coordinates (T1, C1-D1, and D2 in Fig. 1b). To discover neo-TADs and neo-loops from the SV-induced Hi-C contacts, we applied a contig-specific normalization on the contig Hi-C matrices, where the scattered Hi-C contacts are assembled in contig coordinates (contig 1, 2, and 3 in Fig. 1c). Hi-C normalization methods^12,13^ can be applied to the contig Hi-C matrices without the loss of SV-induced Hi-C contacts. Finally, InfoHiC annotates neo-TADs and neo-loops resulting from the SVs on the normalized contig Hi-C matrices (Fig. 1c).

### InfoHiC outperforms the reference-based CNN model

We trained InfoHiC using Hi-C datasets from a breast cancer cell line (T47D) and performed internal validation using it. To evaluate the prediction performance for regions with and without SVs separately, we first extracted genomic regions without SVs. Then, for these regions, we performed five random splits of 4-Mb reference genomic windows, and 80%, 10%, and 10% of regions were used for the training set, validation set, and test set 1, respectively. The remaining genomic regions with SVs, named test set 2, were used for the SV prediction of 4-Mb reference windows and SV windows (novel windows derived from SVs). We measured the distance-stratified correlation (DSC)^9^ and Pearson correlation for reference windows (4-Mb windows that exist in the reference coordinate) of test sets 1 and 2 and measured the Pearson correlation for SV windows of the test set 2. In addition, we used four breast cancer cell lines (MCF7, BT474, SKBR3, and HCC1954) as the external test sets. Test sets were defined for each of these cell lines. Test set_ext_ 1 refers to reference window regions used in test set 1 for T47D after extracting regions with SVs for each cell line and test set_ext_ 2 refers to regions with SVs for each cell line. To evaluate the performance of InfoHiC when non-cancerous cell lines were used for training, we further trained InfoHiC using two breast epithelial cell lines (HMEC and MCF10A), and then tested the four breast cancer cell lines, where the same strategy of five random splits as the T47D was used.

In Fig. 2, the internal validation of the test set 1 (Fig. 2 and Supplementary Table 1) shows performance improvement due to the training strategies and architectures of InfoHiC. First, deepC does not have prior knowledge of SV breakpoints. Thus, we used reference windows around SVs regions (test set 2) for training as well, and obtained the DSC value of 0.607 for the test set 1. When windows with SV-derived contacts were removed for training deepC (breakpoint removal), the performance increased (0.622 DSC). Moreover, HSCN encoding (0.668 DSC) employed in InfoHiC with breakpoint removal showed better performance than HSCN decoding (0.647 DSC) (see Methods), indicating that InfoHiC can learn the relationship between CNAs and Hi-C contact intensities. We then extended the 1-Mb prediction window to 2 Mb to identify TADs greater than 1 Mb. We used the HSCN encoding model for 2-Mb prediction, and the 2-Mb model showed the best performance when the 1-Mb model was transferred (InfoHiC transfer, 0.691 DSC). Finally, we tested the 2-Mb HSCN encoding model with 1-Mb transfer using the external validation of test set_ext_ 1, and showed that our model trained by T47D also predicted external cancer Hi-C data well (InfoHiC transfer, average 0.631 DSC; deepC, average 0.572 DSC), thereby demonstrating the generality of our model.

**Fig. 2:**
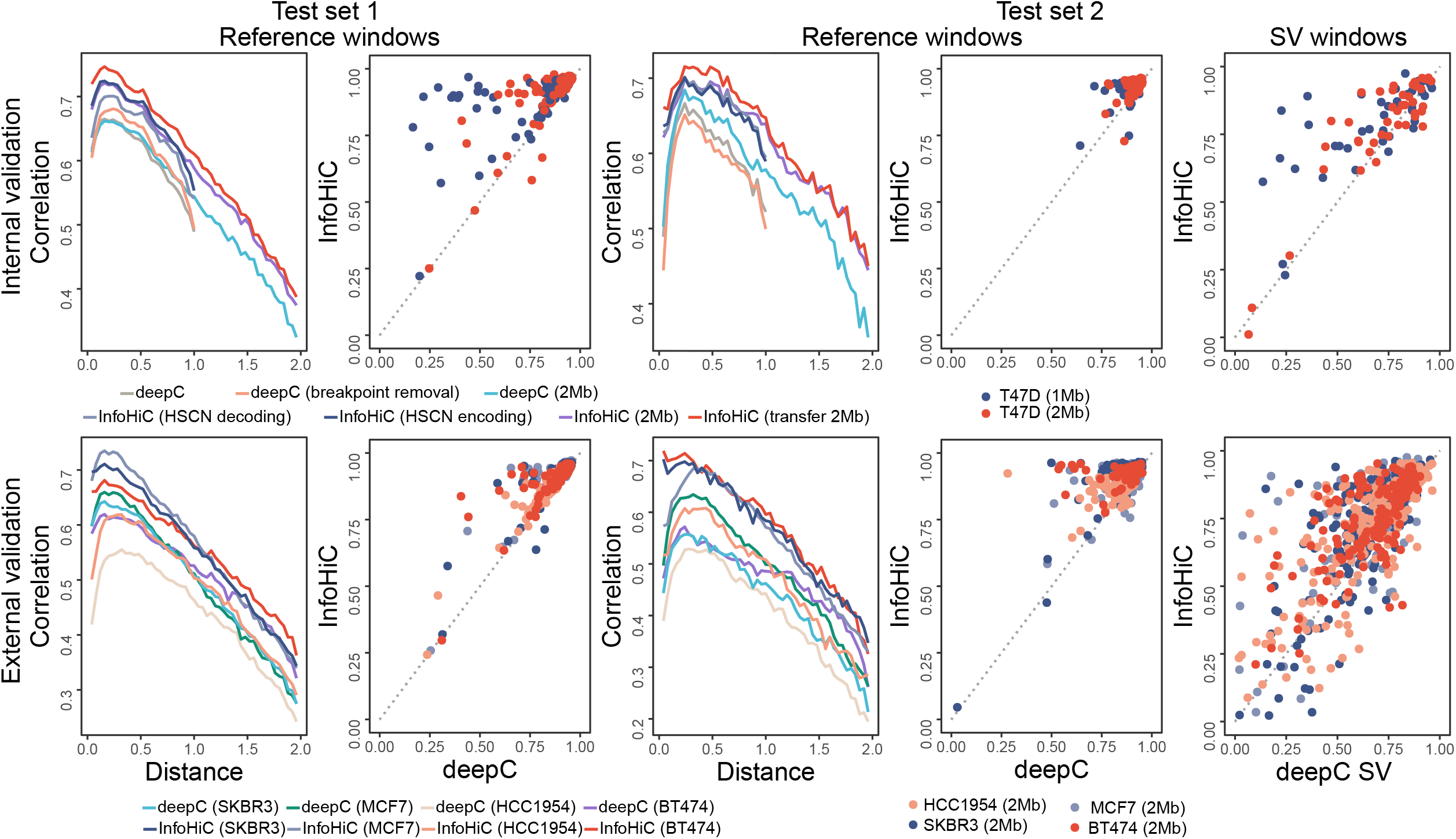
Performance comparison between InfoHiC and deepC. InfoHiC and deepC were compared for training usage of the T47D cell line in test sets 1 and 2. Methods were validated using an internal test set (T47D) at the top and external test sets (SKBR3, MCF7, HCC1954, and BT474) at the bottom. Distance-stratified correlations (line) are shown according to the mega-base scale distance range (0-2 Mb), and the Pearson correlation (dot) for each genomic window was compared between deepC or deepC-SV (bottom) and InfoHiC (left).

Next, we validated InfoHiC for test set 2 containing SV-derived Hi-C contacts (Fig. 2 and Supplementary Table 2), which represent major challenges for cancer Hi-C prediction. For the SV window, deepC-SV, which performed reference training and Hi-C prediction of individual SV calls from InfoGenomeR, was compared with InfoHiC, which performed HSCN training and Hi-C prediction of assembled contigs. InfoHiC (1 Mb, 0.765; 2 Mb, 0.754 Pearson’s R) outperformed deepC-SV (1 Mb, 0.649; 2 Mb, 0.702 Pearson’s R) for T47D (SV window in Fig. 2). The performance was still valid for test set_ext_ 2 (2 Mb InfoHiC, 0.715; 2 Mb deepC, 0.642 average Pearson’s R), demonstrating the predictive power of InfoHiC for SV-induced contacts.

Further, we compared InfoHiC, deepC, and deepC-SV by training using non-cancerous breast epithelial cell lines (HMEC and MCF10A), where performances of deepC and deepC-SV were measured before and after implicit normalization of raw Hi-C data (Supplementary Table 2). For training using HMEC and MCF10A, InfoHiC performed reference training of raw Hi-C data and Hi-C prediction of assembled contigs using HSCN decoding, as HSCN encoding could not be trained by non-cancerous cell lines. InfoHiC showed better performance when the models were trained by HMEC and MCF10A, despite deepC and deepC-SV taking advantage of implicit normalization (Supplementary Fig. 1 and Supplementary Table 2). Validation using external test sets demonstrated that the T47D-trained InfoHiC model showed slightly better performance than the HMEC and MCF10A-trained models, which suggests that training using cancer Hi-C data may be more effective than that using normal-tissue Hi-C data for external cancer Hi-C prediction, or that the Hi-C data of T47D were generated at a higher quality. Under both circumstances, the accumulated cancer Hi-C data would provide benefits to InfoHiC.

### InfoHiC predicts neo-TADs and neo-loops of cancer cell lines

We applied InfoHiC to five breast cancer cell lines and predicted the contig Hi-C matrices. We then performed contig-specific normalization to each contig Hi-C matrix and annotated the neo-TADs and neo-loops. The number of neo-TADs ranged from 47 to 303 (Fig. 3a). More than 20% of neo-TADs were resulted from multiple SVs. In those cases, a Hi-C contact in the cancer Hi-C matrix was not unique to a single TAD, and neo-TADs were overlapped with Hi-C contacts from other TADs (the average number of overlapping TADs was four, with a cut-off of 10% overlapping contacts). These results suggest that assemblies of complex SVs and multiple contig predictions are important for neo-TAD analysis. After annotating typical-enhancer (TE) and super-enhancer (SE) hijacking events of genes in neo-TADs, we searched enhancer hijacking events of several cancer-related genes^14,15^ in the breast cancer cell lines (Supplementary Table 3). InfoHiC can predict SE hijacking events of cancer-related genes from complex SVs such as *MYC* and *PVT1* in SKBR3, which play an oncogenic role in various cancers^16^, as well as *GNA13* in BT474, which is upregulated and associated with poor outcomes in breast cancers^17^. SE hijacking events were confirmed by Hi-C experiments using the InfoHiC validation scheme (Supplementary Fig. 2).

**Fig. 3:**
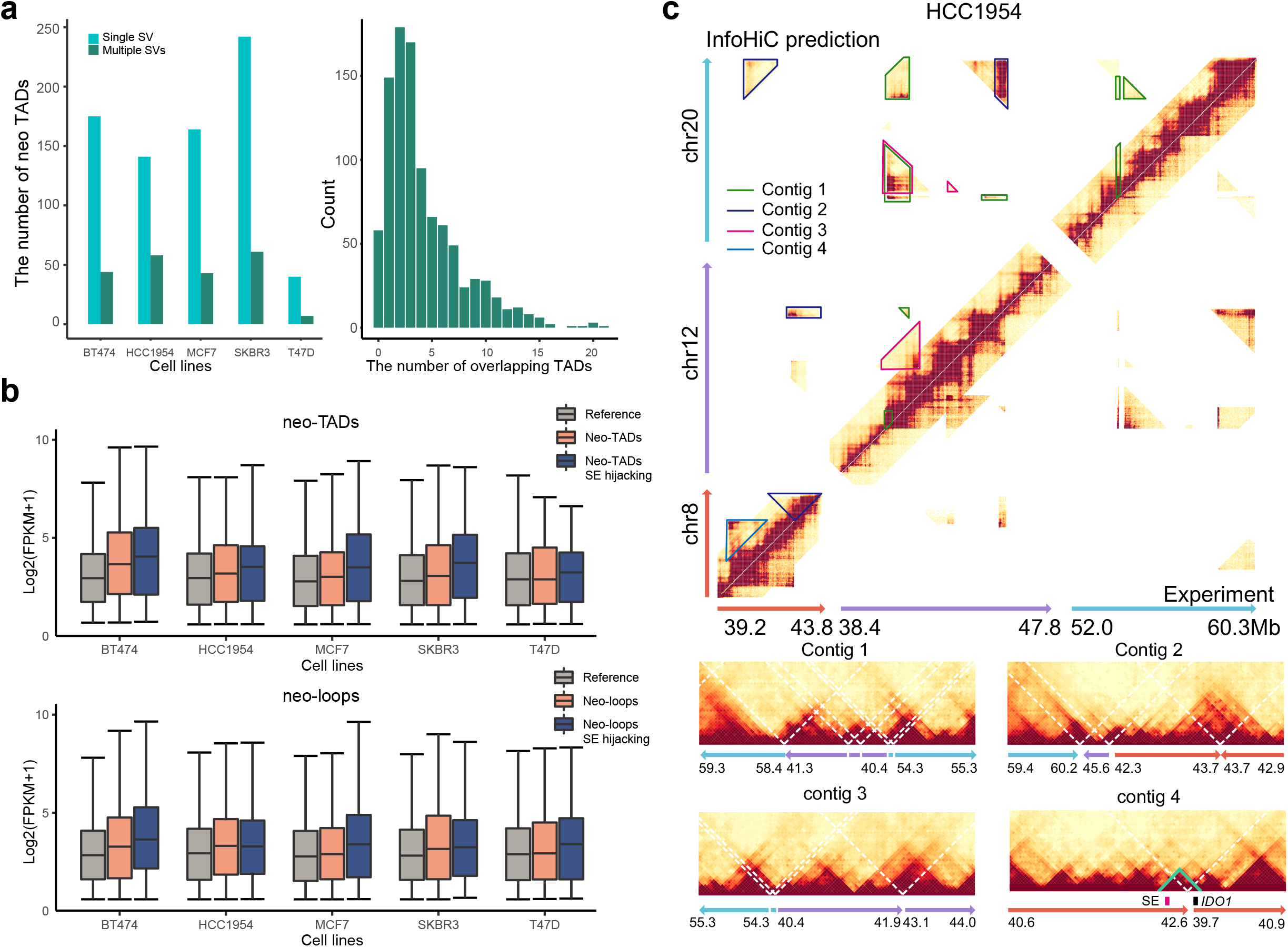
InfoHiC analysis of breast cancer cell lines. **a**, Statistics of the number of neo-TADs in each sample and a histogram of overlapping TAD counts of neo-TADs. **b**, Boxplots of RNA-seq Log2(FPKM+1) values depending on neo-TAD classes (top) and neo-loop classes (bottom) in each sample. Boxplot center lines are medians, box limits are upper and lower quartiles, and whiskers are 1.5x interquartile ranges. **c**, InfoHiC prediction of chromosomes 8, 12, and 20 in the HCC1954 cell line. Hi-C contacts from different genomic contigs (contig 1, 2, 3, and 4) were annotated in the upper diagonal matrix of the total Hi-C (InfoHiC prediction). Lower diagonal matrix represents the Hi-C experiment for HCC1954. Contig Hi-C matrices of each contig are shown below with reference coordinates at the mega-base scale. Dotted white lines represent SV breakpoints. Contig 4 has a neo-TAD (green), where the hijacked SE (crimson) and the *IDO1* gene (black) interact with each other.

Next, we investigated the expression levels of genes in the neo-TADs and neo-loops (Fig. 3b). Genes in neo-TADs were overexpressed compared to those in the reference TADs (Benjamini-Hochberg (BH) adjusted P *<* 0.0001, Student’s one-sided *t*-test), suggesting that neo-TAD formation may induce interactions with other TEs or change the reference chromatin state^4^. Moreover, SE hijacking events resulted in significant overexpression of genes compared to that in the reference TADs (BH-adjusted P *<* 0.0001) and neo-TADs (BH-adjusted P = 0.011). This overexpression was also observed in neo-loop annotations (BH-adjusted P *<* 0.0001 in all three comparisons). Furthermore, we performed analysis of covariance (ANCOVA) by controlling for CN effects on gene expression (Supplementary Fig. 3). Overexpression by neo-TADs and SE hijacking events were observed under CN covariates (P = 0.001, one-way ANCOVA), suggesting that neo-TADs and SE hijacking events are involved in the dysregulation of gene expression in cancer.

An example of neo-TAD formation in the *IDO1* gene was observed in the HCC1954 cell line. HCC1954 had three distinct SV clusters, one of which was found among chromosomes 8, 12, and 20 (Fig. 3c). Moreover, SV clusters resulted in neo-TADs from different contigs. A tandem duplication was found near *IDO1*, which resulted in an SE hijacking event (contig 4 in Fig. 3c). Interaction between *IDO1* and the SE was absent in the other breast cell lines, indicating a neo-interaction. The *IDO1* gene showed the highest fragments per kilobase of transcripts per million mapped reads (FPKM) level among breast cancer cell lines from the Cancer Cell Line Encyclopedia (CCLE) (Supplementary Fig. 4), whereas integer CNs of the genes did not significantly affect gene expression (−0.002 Pearson’s R). The correlation between *IDO1* expression and the intracellular level of an immunosuppressive metabolite, kynurenine, was reported in a recent CCLE report^18^. The HCC1954 cell line showed the second-highest kynurenine levels among the breast cancer cell lines. Collectively, these results showed that InfoHiC can provide evidence of the 3D genome context associated with *IDO1* overexpression and high kynurenine levels.

**Fig. 4:**
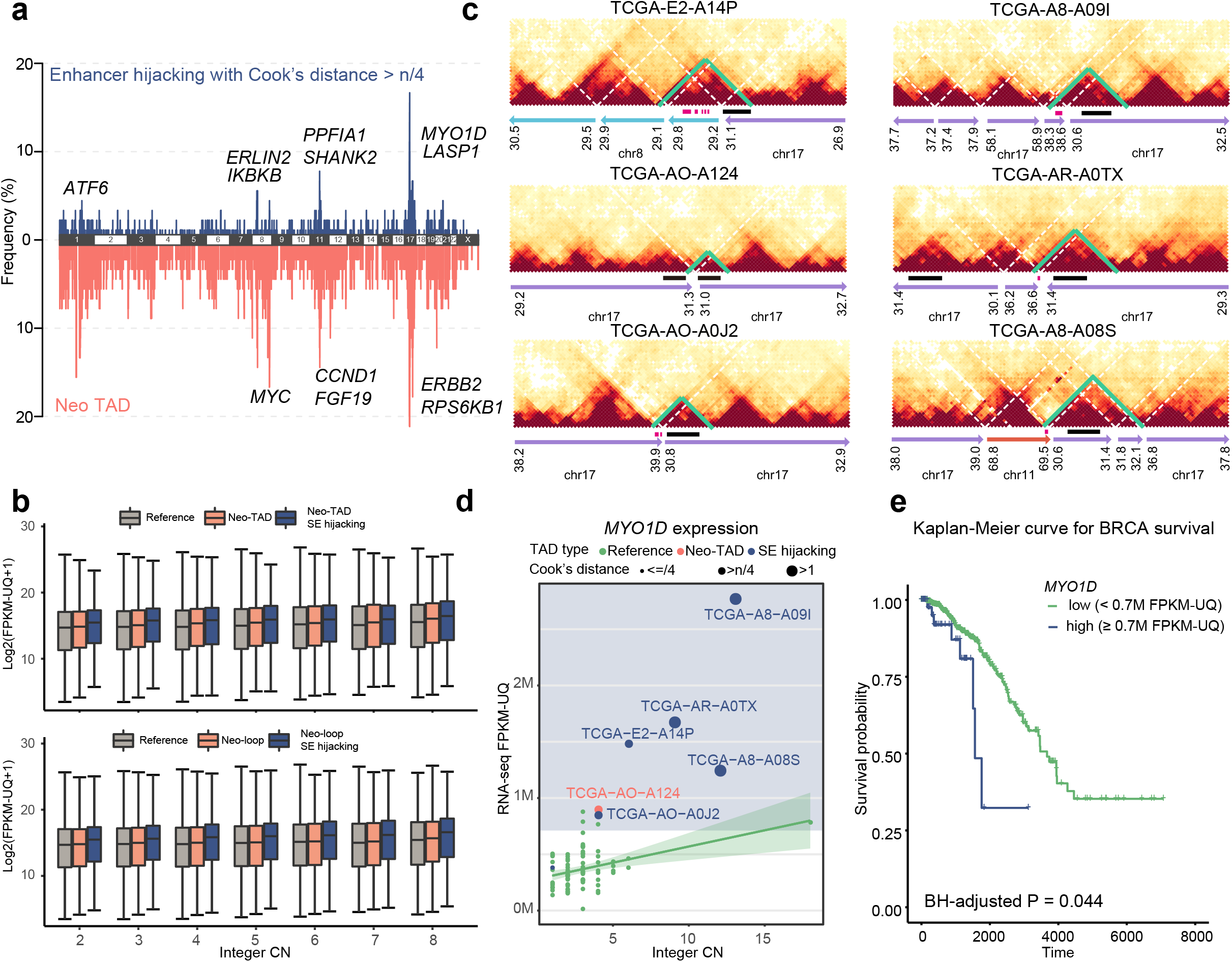
InfoHiC analysis of patients with TCGA BRCA. **a**, Frequencies of neo-TADs (apricot) and enhancer hijacking events (blue) associated with overexpression across chromosomes. Each peak represents the neo-TAD or enhancer hijacking frequency of each gene. The representative peaks were annotated using gene symbols. **b**, Boxplots of RNA-seq Log2(FPKM-UQ+1) values depending on neo-TAD classes (top) and neo-loop classes (bottom) in patients with BRCA according to integer CNs. Boxplot center lines are medians, box limits are upper and lower quartiles, and whiskers are 1.5x interquartile ranges. **c**, InfoHiC prediction of contig Hi-C matrices of patients with BRCA. Neo-TADs are shown in green, with hijacked SEs (crimson) and the *MYO1D* gene (black). **d**, Gene expression of *MYO1D* of patients with BRCA (dot) using WGS data (n=90). A linear regression line between FPKM-UQ values and integer CNs is shown with the corresponding confidence interval range (green). Neo-TAD (apricot) and SE hijacking samples (blue) are shown in different circle sizes according to the Cook’s distance. Cut-off region of the FPKM-UQ value for overexpression is shown in the background (blue). **e**, Kaplan–Meier curve for BRCA survival with RNA-seq data (n=1,098). P value was calculated using the log-rank test and adjusted using the BH procedure across neo-TAD overexpression genes.

### Non-coding effect of SVs in 3D cancer genomes

Subsequently, we applied InfoHiC to BRCA patients (n=90) in TCGA to investigate the non-coding effects of SVs. Neo-TADs were annotated on contig Hi-C prediction and were found with high frequency peaks in chromosomes 1, 8, 11, 17, and 20 (bottom plots in Fig. 4a). Recurrent neo-TAD genes (found in *≥* 10% neo-TAD samples) were enriched in the KEGG breast cancer pathway (BH-adjusted P = 0.014), including breast cancer genes such as *FGF19, ERBB2, CCND1, MYC*, and *RPS6KB1*. Genes in neo-TADs were overexpressed under CN covariates (P *<* 0.0001, one-way ANCOVA), demonstrating the non-coding effects of SVs on gene expression (Fig. 4b).

We also observed that neo-TAD formation frequently accompanies CNAs in breast cancer. To identify recurrent neo-TAD genes associated with overexpression controlling for CNAs, we performed a linear regression between gene CNs and expression in reference TAD samples without neo-TADs. Based on the regression line, we measured Cook’s distance (an indicator of outlier influence on the regression line)^19^ for the neo-TAD samples. We counted the number of neo-TAD samples with a positive residual and a Cook’s distance greater than 4/n (n=90), which indicated overexpression compared with the reference TAD samples. Genes with such neo-TADs were found to have peaks in the 8p11, 11q13, and 17q11 regions (top plots in Fig. 4a). We selected genes with the following attributes: 1) more than 50% of neo-TAD samples showing a Cook’s distance greater than 4/n and 2) with SE hijacking samples *≥* 4 as recurrent neo-TAD genes involved in overexpression (Supplementary Table 4), which included known driver genes^14^ such as *ERBB2, CCND1, LASP1, CLTC*, and *CDK12*. We further investigated the survival outcomes of patients with recurrent neo-TAD genes across all BRCA data using RNA-seq from TCGA (n=1098). A threshold for overexpression samples was defined as the minimum FPKM value over the Cook’s distance cut-off for each gene (Supplementary Table 4). We found three recurrent neo-TAD genes (*LASP1, CLTC*, and *MYO1D*) with poor prognosis. *LASP1* overexpression is reportedly correlated with poor prognosis in breast cancers; however, the cause of *LASP1* overexpression remains unclear (it does not result from CNAs)^20^. Here, we discovered neo-TAD formation of *LASP1* associated with overexpression and poor prognosis (BH-adjusted P = 0.019, log-rank test) (Supplementary Fig. 5). The most recurrent neo-TAD formation associated with poor survival was found in the *MYO1D* gene (Fig. 4c). *MYO1D* is in a repressed TAD, indicating that its transcriptional activity is low in the reference state^4^. Seven patients exhibited neo-TAD formation of *MYO1D*. We found SE hijacking events with neo-TAD formation in six patients, five of whom showed overexpression of the *MYO1D* gene (Fig. 4d). One patient (TCGA-AO-A124) did not have a SE in the neo-TAD but also showed overexpression of the *MYO1D* gene, as tandem duplication may have a position effect on gene expression^21^. Furthermore, overexpression of *MYO1D* was a prognostic factor for survival in TCGA BRCA patients (BH-adjusted P = 0.044, log-rank test) (Fig. 4e). Overexpression of *MYO1D* has previously been associated with breast cancer cell motility and viability^22^. Collectively, these results suggest that InfoHiC can detect 3D genome changes associated with overexpression and poor prognosis.

**Fig. 5:**
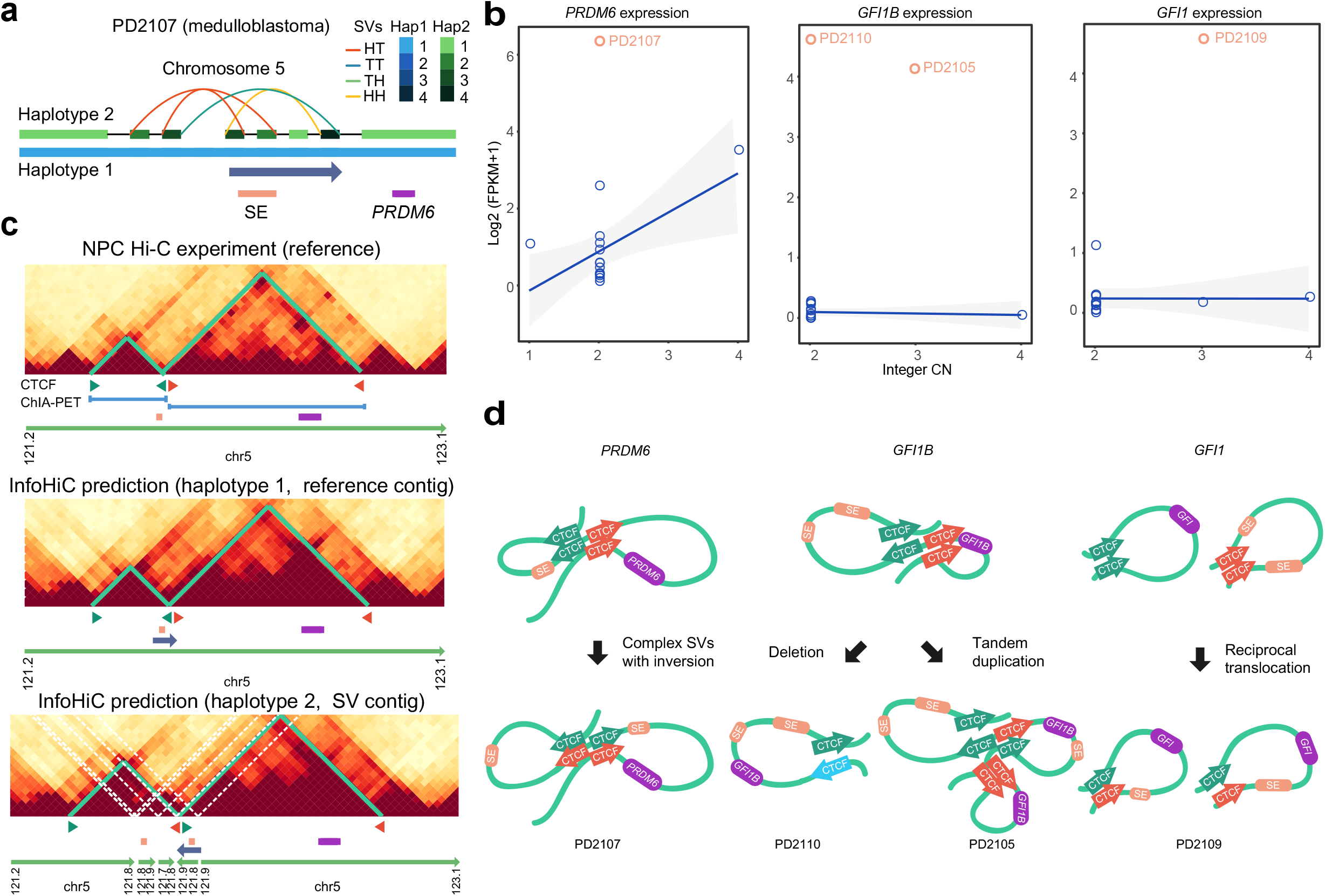
Super-enhancer (SE) hijacking events of *PRDM6, GFI1B*, and *GFI1* found in patients with medulloblastoma. **a**, A haplotype graph of chromosome 5 of the patient with medulloblastoma (PD2107). SVs were annotated with head-to-tail (HT), tail-to-head (TH), tail-to-tail (TT), and head-to-head (HH) orientations. **b**, FPKM values of the *PRDM6, GFI1B*, and *GFI1* transcripts versus integer CNs measured in the brain tumor cohort. Linear regression lines are shown with the corresponding confidence interval ranges (blue), and the FPKM values of patients (PD2105, PD2107, PD2109, and PD2110) with SE hijacking events are shown above (apricot). **c**, Hi-C experiment of neuro-progenitor cells (NPCs) and InfoHiC prediction of the 3D genome of patient PD2107. Reference TADs and neo-TADs are annotated (green) with CTCF motifs (arrowheads) near the boundaries, and *PRDM6* (purple) and SE (apricot) are shown below. The pairwise interactions of the CTCF ChIA-PET data are indicated by blue lines. **d**, 3D genome models of the SE hijacking events of *PRDM6, GFI1B*, and *GFI1* derived from various types of SVs.

### Application of InfoHiC to pediatric patients with medulloblastoma

To demonstrate the analytical potential of InfoHiC in discovering non-coding driver SVs, we applied InfoHiC to five pediatric patients with medulloblastoma in our cohort. After we performed WGS and RNA-seq analysis to these patients, the presence of driver mutations in coding regions were examined using cancer-related gene lists^14,15,23^ (Methods). Driver mutations were not found in three patients (PD2104, 2107, and 2109) by single nucleotide variants (SNVs), indels, and CNA calls (Supplementary Table 5 and 6). *MYC* amplification, which is a prevalent driver of group 3 medulloblastomas^24^, and LOH deletion of a tumor suppressor, *TSC1* was found in the PD2105 and PD2110 patients, respectively. Further analysis of SVs using InfoGenomeR revealed derivative chromosomes in the medulloblastoma patients; however the impacts of SVs were unknown as most SVs occured in non-coding regions and were not relevant to CN changes or fusion events of driver genes (Supplementary Fig. 6-9). Next, we investigated SVs that may affect the 3D genome organization. Notably, we found complex SVs in the non-coding region in the driver mutation-negative case (PD2107), which included a subtype-specific SE of group 4 medulloblastomas near the *PRDM6* gene, the overexpression of which has been suggested as a driver of medulloblastoma^23^(Fig. 5a). The patient also showed overexpression of *PRDM6* with a 90-fold change in FPKM compared to that in our cohort of brain tumors, suggesting that the complex SVs may induce *PRDM6* over-expression (Fig. 5b). In addition, we found recurrent SVs near the *GFI1* family oncogenes; a deletion and tandem duplication near the *GFI1B* gene in the *TSC1*-deleted and *MYC*-amplified cases (PD2110 and PD2105, respectively), and reciprocal translocation near the *GFI1* gene in the driver mutation-negative case (IITP2109). The patients showed overexpression of *GFI1* family oncogenes with more than 100-fold changes in FPKM (Fig. 5b), suggesting that these SVs could be medulloblastoma drivers associated with oncogenic overexpression.

Using InfoHiC, we discovered neo-TAD formation derived from complex SVs and a SE hijacking event in *PRDM6* (Fig. 5c). We then annotated the CTCF motif directions in CTCF ChIP-seq peak regions of TAD boundaries using PWMScan^25^ and checked the pairwise interactions of the CTCF ChIA-PET data^26^, which supported the hypothesis that the reference TADs were arranged in forward-reverse orientations. Complex SVs included an inversion spanning from the SE to the CTCF motifs, which resulted in the SE hijacking event of the TAD containing *PRDM6* (Fig. 5c). This inversion maintained the forward-reverse CTCF arrangements, supporting neo-TAD formation with the SE (Fig. 5d). Complex SVs in the non-coding region near *PRDM6* were also reported in group 4 medulloblastomas in a previous study^23^, whereas neo-TAD formation by complex SVs remains unclear. Our results demonstrated that InfoHiC can validate neo-TAD formation by complex SVs and specify the non-coding driver SVs from the driver mutation-negative case (PD2107). Furthemore, InfoHiC revealed SE hijacking events of neo-TADs of *GFI1* oncogene families induced by various types of SVs (PD2105, 2109, and 2110) (Supplementary Fig. 10 and Fig. 5d); a deletion occured across TAD boundaries causing TAD fusion (PD2110), a tandem duplication spanned an SE and the *GFI1B* gene causing a neo-TAD with the hijacked SE (PD2105), and a reciprocal t(1;5) translocation caused interchromosomal neo-TADs with distant SEs (PD2109), demonstrating that neo-TADs were involved with activation of *GFI1* oncogene families.

## Discussion and Conclusions

In summary, we developed InfoHiC for cancer Hi-C prediction using a sequence-based model that enables the identification of neo-TAD and neo-loop formation from rearranged genomes. In addition to providing a validation scheme for SV-derived Hi-C contacts, InfoHiC performs better than the reference-based model. Currently, there is no normalization method suitable for the cancer Hi-C matrix dealing with SV-derived contacts from multiple contigs. Therefore, instead of applying normalization methods to cancer Hi-C data to remove CNAs and SVs as biases, InfoHiC uses raw Hi-C data for training and performs contig-specific normalization after contig Hi-C prediction. Thus, InfoHiC provides normalized contig Hi-C matrices, which can be analyzed using other analytics tools developed for Hi-C data.

Regarding previous studies aimed at obtaining a contig Hi-C matrix from cancer Hi-C data, NeoLoopFinder^8^ uses one-to-one mapping from the reference coordinate to a single contig coordinate, which restricts cancer Hi-C data to large SVs (*>*1 Mb). However, InfoHiC uses a many-to-one mapping from contig coordinates to the reference coordinate; thus InfoHiC is available for any type of SVs, as shown in the tandem duplication, deletion, translocations, and cluster of complex SVs (*<* 1 Mb) in the medulloblastoma cases (Fig. 5). Compared with reference-based prediction models such as deepC^9^, Akita^10^, and Orca^11^, InfoHiC enables the use of cancer Hi-C data for training and validation, which takes advantage of the accumulated Hi-C datasets of cancer cell lines. In addition, applications of reference-based models^9–11^ are currently limited to simple deletions, duplications, or inversions; however, we showed that Hi-C prediction from complex SVs is essential for neo-TAD findings in cancers.

In the application study of pediatric patients with medulloblastoma, we showed that various types of SVs caused neo-TAD formation of the same *GFI1* families. In addition, the impact of complex SVs was uncovered by predicting Hi-C interaction from the assembled contig. Previous approaches of SV analysis in non-coding regions include annotating juxtaposition events between genes and regulatory elements^24^ or boundary-affecting SVs from known TAD boundaries^4^. However, SV analysis requires Hi-C evidences as chromatin interaction between DNA elements and TAD boundaries change according to rearranged sequence contexts. InfoHiC annotates juxtaposition events (SE hijacking) and neo-TAD boundaries based on Hi-C prediction, providing more accurate results for analyzing non-coding SVs. Furthermore, the impact of complex SVs could not be investigated by previous approaches of enhancer-juxtaposing^24^ or boundary-affecting SV annotation, and InfoHiC provides Hi-C evidences for neo-TAD boundaries derived from complex SVs. Based on InfoHiC prediction, targeted therapy may be possible for non-coding driver SVs. For the example of medulloblastoma (PD2109) where GFI1 gene was overexpressed by the reciprocal translocation t(1;5), inhibitors of a co-factor of *GFI1* transcription factor, *LSD1* can be used for targeted treatment. *LSD1* inhibitors such as GSK-LSD1 and ORY-1001 were shown to be effective for *GFI1*-activated medulloblastoma^27^.

As an integrative work from 1D genome reconstruction to 3D genome prediction, we demonstrated that SVs have non-coding effects on the overexpression of cancer-related genes, which are associated with poor prognosis in BRCAs. We expect that the application of InfoHiC to other cancer types may improve the chances of identifying cancer drivers altered by non-coding SVs, especially for cancer types without driver SNVs and CNA events in coding regions. Furthermore, future research may identify cancer drivers in patients without common coding driver mutations, which may lead to personalized medication based on patient-specific SVs in the 3D genome context.

## Methods

### Reconstruction of genomic contigs

Genomic contigs were obtained using InfoGenomeR from WGS data, which were used to construct a breakpoint graph to model the connectivity among genomic segments using SVs, CNAs, and SNP information and derived genomic contigs from Eulerian paths according to a minimum entropy approach^1^. We used DELLY2^28^, Manta^29^, and novo-Break^30^ for the initial SV detection from Illumina paired-end data. Then, SV calls pre-detected from other WGS platforms (see Data availability section) were merged into the initial SV detection. We used BIC-seq2^31^ for CNA detection and ABSOLUTE^32^ for integer-CN detection (with cancer purity and ploidy estimation). SAMtools and BCFtools^33^ were used for SNP detection, and the haplotype-cluster model of BEAGLE^34^ from 1000G was used for haplotype phasing in InfoGenomeR.

### HSCN encoding

The genomic contigs were derived from SVs in haplotypes, and had different CNs and SNP compositions with one another. Their sequences were represented by HSCN encoding, which was modified from one-hot encoding to represent the CNs and SNPs in a contig matrix. The column of the contig matrix represents each base index *j* of the contig, whereas the row index *i* represents the base composition, i.e., A, C, G, and T. One-hot encoding represents a zero or one binary for the base, and HSCN encoding multiplies the binary by the CN of the contig. We denote the genome *G* as a set of genomic contigs *g*, with length *l* and HSCN encoding function as *H*. The CN of the genomic contig was *µ*(*g*).

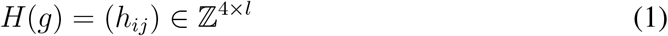

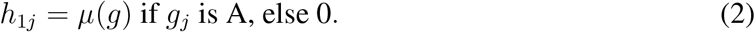

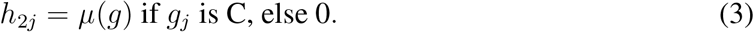

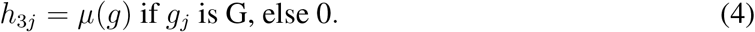

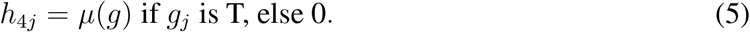

### Hi-C data processing

We used the 3DIV pipeline for Hi-C read mapping, which previously established the first large-scale resource for cancer Hi-C^35^. Reads were mapped to the hg19 reference genome. The chimeric and self-ligated reads were filtered using the 3DIV pipeline. The contact matrix was obtained at a resolution of 40 kb. In the InfoHiC, we used the raw contact matrix without implicit normalization for training and validation to preserve the Hi-C signals from genomic rearrangements. For comparison, we applied covNorm^13^, a normalization method included in the 3DIV pipeline, to the raw contact matrix for implicit normalization.

### Prediction of the total Hi-C matrix

We denote the convolutional neural network as CNN, which is composed of CNN_deepSEA_ to extract chromatin features from DNA sequences^36^, a dilated convolution neural network CNN_dilated_ to model a wide range of contexts for contact prediction^9^, and a fully connected layer *FC*_*d*_ to predict the Hi-C contact for a genome distance *d*. The target value of the Hi-C interaction is *c*(*x, y*), which represents the number of Hi-C reads mapped into the *x*- and *y*-coordinate bins in the Hi-C matrix. Here, we denote *c*(*x, y*) as *c*. The reference version of deepC^9^ predicts *c* using a single reference contig centered on *c*. In a reference model, the target *c* is in the reference coordinate. We denote one-hot encoding based on the reference sequence as *R*.

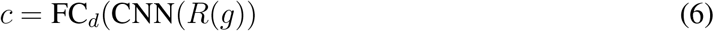

In contrast to the reference model, we predict *c* as the sum of Hi-C contacts of multiple contigs, *g*_1_, *g*_2_, …, *g*_*n*_ with different genomic distances *d*_1_, *d*_2_, … *d*_*n*_ centered on target values *c*_1_, *c*_2_, …, *c*_*n*_ in the contig coordinates. When using different contigs rather than a reference sequence, we require a reference coordinate mapping function to change the contig coordinate prediction, which is denoted by ref.

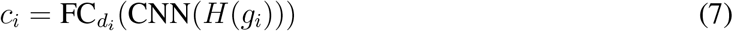

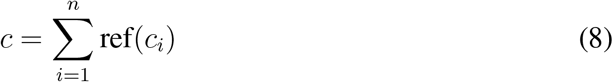

The ref(*c*_*i*_) function indicates the mapping from *c*_*i*_(*x*^*′*^, *y*^*′*^) on the *x*^*′*^ and *y*^*′*^ contig into *c*_*i*_(*x, y*) on the *x* and *y* reference coordinates. It returns *c*_*i*_ if two 40-kb genomic bins (*x*^*′*^ and *y*^*′*^) targeting *c*_*i*_ in the contig coordinate are maximally matched with two genomic bins (*x* and *y*) targeting *c* in the reference coordinate; otherwise, it returns 0. When Hi-C reads were mapped at a certain resolution, a genomic bin in the contig coordinates contained multiple portions of several bins in the reference coordinate. For example, a 40-kb genomic bin in the contig coordinate could contain 30 kb of bin 1 and 10 kb of bin 2 in the reference coordinate. In this case, we select a maximum match (bin 1) to maintain the sharpness of the Hi-C data, which enables accurate TAD annotation. For training multiple contigs with SV, there are issues compared to training the reference sequence. The reference prediction such as deepC^9^ uses a single one-hot encoding matrix to predict a vector of targets *c*=(*c*^1^, *c*^2^, … *c*^*k*^) using a vector of genomic distances *d*=(*d*^1^, *d*^2^, … *d*^*k*^) (a zig-zag pole), simultaneously, rather than targeting a single scalar of *c*^*k*^. However, for multiple contigs with SVs, prediction of the target vector is complicated. First, the genomic distances could differ when targeting *c*^*k*^. Second, the reference coordinate mapping should be included in the training. Third, tens of contigs could exist, and the graphics processing unit (GPU) memory would not be available for them. Thus, to simplify the training procedure for multiple contigs, we used genomic contigs in reference regions without SVs for training (breakpoint removal). The following are true for the reference regions: 1) genomic distances are the same for two haplotypes; 2) the ref function is not required because it is in the reference coordinate itself; 3) only two haplotype contigs are required to enable proper GPU memory usage. Simply, we target a vector of target *c*=(*c*^1^, *c*^2^, …, *c*^*k*^) using two haplotype contigs, *g*_1_ and *g*_2_, with the genomic distance *d*^1^, *d*^2^, …, *d*^*k*^. Here, *c*^*j*^ is the sum of the haplotype target values for each (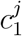 and 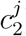).

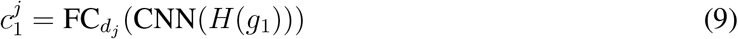

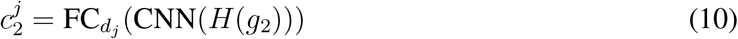

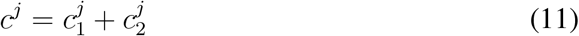

Given the two haplotype contigs for training, we denote *ĉ* = (*ĉ*^1^, *ĉ*^2^, …, *ĉ*^*k*^) as a zig-zag pole prediction^9^ and minimized the mean square error (MSE) loss between *c* and *ĉ*.

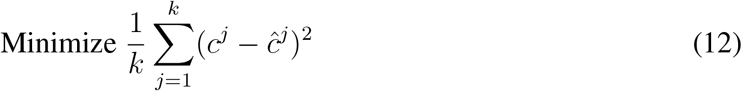

### HSCN decoding

HSCN decoding was used for a comparison study with HSCN encoding when InfoHiC was trained by the cancer cell line (T47D). In addition, when InfoHiC was trained by the non-cancerous cell lines (HMEC and MCF10A), we used HSCN decoding for the InfoHiC prediction model because HSCN encoding could not be trained by these cell lines. In HSCN decoding, haplotype contigs, *g*_1_, *g*_2_, …, *g*_*n*_ are encoded by the one hot encodings of haplotype bases, *H*_one-hot_(*g*_*i*_), *H*_one-hot_(*g*_*i*+1_), …, *H*_one-hot_(*g*_*n*_), and prediction outputs are multiplied by the CN of each genomic contig.

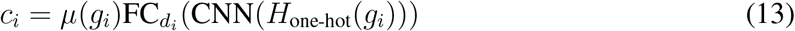

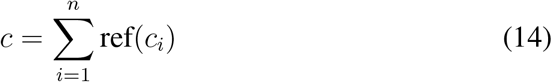

### CNN architecture

We used a deepC architecture^9^ to predict Hi-C contacts per genome contig. DeepC was previously adapted from DeepSEA^36^ that employs hundreds of convolution filters to extract chromatin features. Specifically, five convolution layers (300, 600, 600, 900, and 900 hidden units) with various kernel widths (8, 8, 8, 4, and 4) and max-pooling schemes (4, 5, 5, 5, and 2) were used to extract the chromatin features. The layers were pretrained using chromatin profiling data (936 chromatin features). Then, 10 dilated gated convolutional layers (100 hidden units) with various dilation schemes (1, 2, 4, 8, 16, 32, 64, 128, 256, and 1) were used for long-range modeling, and the final fully connected layer output the contact value for each distance.

### Transfer learning and data augmentation

We used two transfer-learning steps in this study. First, we used the pretrained weights of CNN_deepSEA_ from DeepC, which was trained using input 1-kb DNA sequences targeting 936 chromatin features. We then transferred the pretrained weights of CNN_deepSEA_ and trained the convolutional neural network CNN = CNN_dilated_ ° CNN_deepSEA_ for the total Hi-C matrix prediction using 1-Mb + 40-kb genomic windows at a resolution of 40 kb, targeting a vector of 25 contact values, *c* = (*c*_1_, *c*_2_, …*c*_25_). Then, we transferred the trained weights from 1-Mb + 40-kb genomic windows to the 2-Mb prediction model, targeting a vector of 49 contact values, *c* = (*c*_1_, *c*_2_, …*c*_49_). We excluded self-interaction (zero distance interaction in a single 40-kb bin) because raw Hi-C data have high values in self-interaction that prevented the learning of long range interactions.

We then used augmentation processes including a 20-kb shift and reverse complement training. For the 20-kb shift, we mapped the Hi-C reads to +20-kb genomic bins. In detail, Hi-C reads were mapped to the 1-kb to 40-kb coordinate in the first mapping process, and Hi-C reads were mapped to the 21-kb to 60-kb coordinate in the second mapping, producing a doubled training data set. We trained the model using the forward strand for the original Hi-C and the reverse strand for the flipped Hi-C. For testing, we averaged the predictions of the forward and reverse strands.

### Contig-specific normalization

Normalization methods commonly used for the Hi-C matrix, such as the iterative correction and eigenvector decomposition (ICE)^12^, consider CNAs and SVs as biases. ICE normalization was employed to implicitly normalize the GC content bias, mappability bias, and other experimental noise together with CNA and SV biases by introducing biases *b*_*i*_ and *b*_*j*_ for each *i*th and *j*th bin. Here, *c*(*i, j*) is an observation of the total Hi-C contact, *t*(*i, j*) is the normalized Hi-C contact, and *N* is the number of bins.

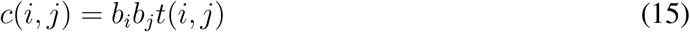

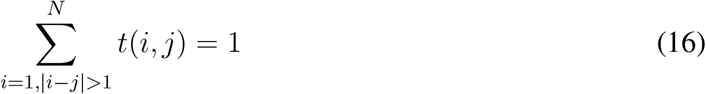

When using the total Hi-C observation *c*(*i, j*) for cancer Hi-C, it can result in inaccurate production of a normalized matrix for the reference sequence. In addition, Hi-C contacts from CNAs and SVs should not be removed to discover 3D genome organization for rearranged contigs. Previously, we obtained *ĉ* for *n* assembled contigs.

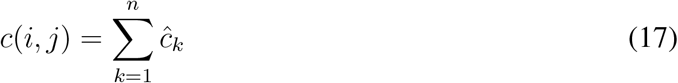

Here, we applied ICE normalization to *ĉ*_*k*_, which is an assembled Hi-C contact free from CNA and SV biases. Here, *i*^*′*^ and *j*^*′*^ are indices in contig coordinates for each *k* contig. The Iced Python module was used for contig-specific ICE normalization.

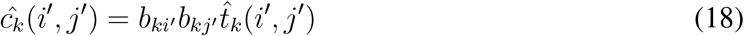

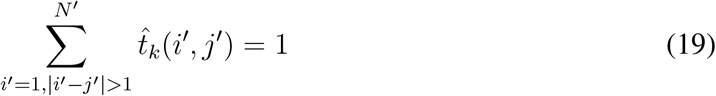

### Neo-TAD and neo-loop annotation

We annotated the TADs and loops on the normalized contig Hi-C matrix 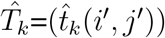 using spectralTAD^37^ and Peakachu^38^, respectively. We hierarchically obtained primary to tertiary TADs using spectral clustering^37^. Neo-loops were annotated using the CNN-based method Peakachu^38^ on the contig Hi-C obtained by InfoHiC, which is free of CNA and SV biases and does not require any further steps of CN normalization or pseudo-assembly of SV-derived contacts^8^. A TE or SE hijacking event of a gene in the neo-TAD was defined as follows: an enhancer did not exist in the reference TAD prediction (with a *±*40-kb offset to adjust errors of TAD boundaries) and 2) interaction between the enhancer and the gene was predicted to exist in the SV window.

### Sample preparation and data preprocessing

For brain tumors, macrodissection was performed on tumor areas with a tumor cell content of more than 90% stained with hematoxylin-eosin. Genomic DNA and total RNA were extracted from the freshly frozen tissues and peripheral blood of patients using the Maxwell RSC DNA and RNA FFPE Kit (Promega, Madison, WI, USA), respectively. The WGS library was prepared using the TruSeq Nano DNA Kit (Illumina, San Diego, CA, USA) and sequenced using the Illumina NovaSeq6000 platform. The RNA-seq library was prepared using the SureSelectXT RNA Direct Library Kit (Agilent Technologies, Santa Clara, CA, USA) and sequenced using the Illumina NovaSeq6000 platform.

Paired-end WGS reads (100 bp) from the T47D, MCF7, and SKBR3 cell lines and paired-end WGS reads (150 bp) from brain tumors including the medulloblastoma case were mapped to the human reference genome (GRCh37) using BWA-MEM^39^ with default parameters (version 0.7.15). Somatic SNVs and indels were detected using Mutect^40^ and Platypus^41^ respectively, and annotated using ANNOVAR^42^. Paired-end RNA-seq reads (101 bp) from brain tumors were mapped to GRCh37 using STAR^43^, and gene expression was quantified using Cufflinks^44^.

## Data availability

WGS data of TCGA samples are available from dbGaP (accession code phs000178.v11.p8) [https://www.ncbi.nlm.nih.gov/projects/gap/cgi-bin/study.cgi?studyid=phs000178.v11.p8]. RNA-seq gene expression data of TCGA samples were downloaded from the GDC data portal. Hi-C fastq files of breast cancer cell lines are available from the sequencing read archive (SRA); MCF7, BT474, and SKBR3 Hi-C data (accession codes PRJNA430222); HCC1954 (accession code PRJNA479882); and T47D (PRJNA438511), respectively. The WGS fastq files of the T47D, MCF7, and SKBR3 cell lines are available from the SRA: T47D (PRJNA380394), MCF7 (PRJNA486532), and SKBR3 (PRJNA476239). WGS bam files of the HCC1954 and BT474 cell lines were downloaded from TCGA in the GDC data portal and CCLE in Google Cloud Storage, respectively. SV calls of HCC1954 from linked-read sequencing data (10X Genomics) were downloaded from https://cf.10xgenomics.com/samples/genome/HCC1954TWGS210/HCC1954T_WGS_210_large_svs.vcf.gz. SV calls for SKBR3 from long-read sequencing data (PacBio)^45^ were downloaded from http://labshare.cshl.edu/shares/schatzlab/www-data/skbr3/reads_lr_skbr3.fa_ngmlr-0.2.3_mapped.bam.sniffles1kb_auto_l8_s5_noalt.vcf.gz. Hi-C fastq files of NPC cells are available from SRA (accession code PRJNA798046). CTCF ChIP-seq data of NPC cells and ChIA-PET data of a neuroblastoma cell line are available from ENCODE (accession code ENCSR125NBL and ENCSR514HBO, respectively).

## Code availability

InfoHiC is available on GitHub https://github.com/dmcb-gist/InfoHiC.

## Acknowledgements

This work was supported by an Institute of Information & Communications Technology Planning & Evaluation (IITP) grant funded by the Korean government (MSIT) (No. 2019-0-00567, Development of Intelligent SW Systems for Uncovering Genetic Variation and Developing Personalized Medicine for Cancer Patients with Unknown Molecular Genetic Mechanisms). The brain tumor biospecimens and data used in this study were provided by the Human Biobank of Seoul National University Hospital, a member of Korea Biobank Network (KBN4 A03), and Seoul National University Hospital Cancer Tissue Bank. All samples derived from the Biobanks of SNUH were obtained with informed consent under institutional review board-approved protocols.

## Author contributions

HL initiated and supervised the project. HL, YL, and SP collected data and analyzed the results. HL and YL developed the algorithm and wrote the manuscript. YL performed the experiments.

## Ethics declarations

The study on brain tumors was approved by the Institutional Review Board of Seoul National University Hospital (IRB No:1905–108-1035). All experiments were performed in accordance with the guidelines and regulations of the Helsinki and Human Research Protection Programs.

