## Supplementary Figures for "Prediction of the 3D cancer genome from genomic rearrangements using InfoHiC"

### Supplementary Information for Prediction of the 3D cancer genome from genomic rearrangements using InfoHiC

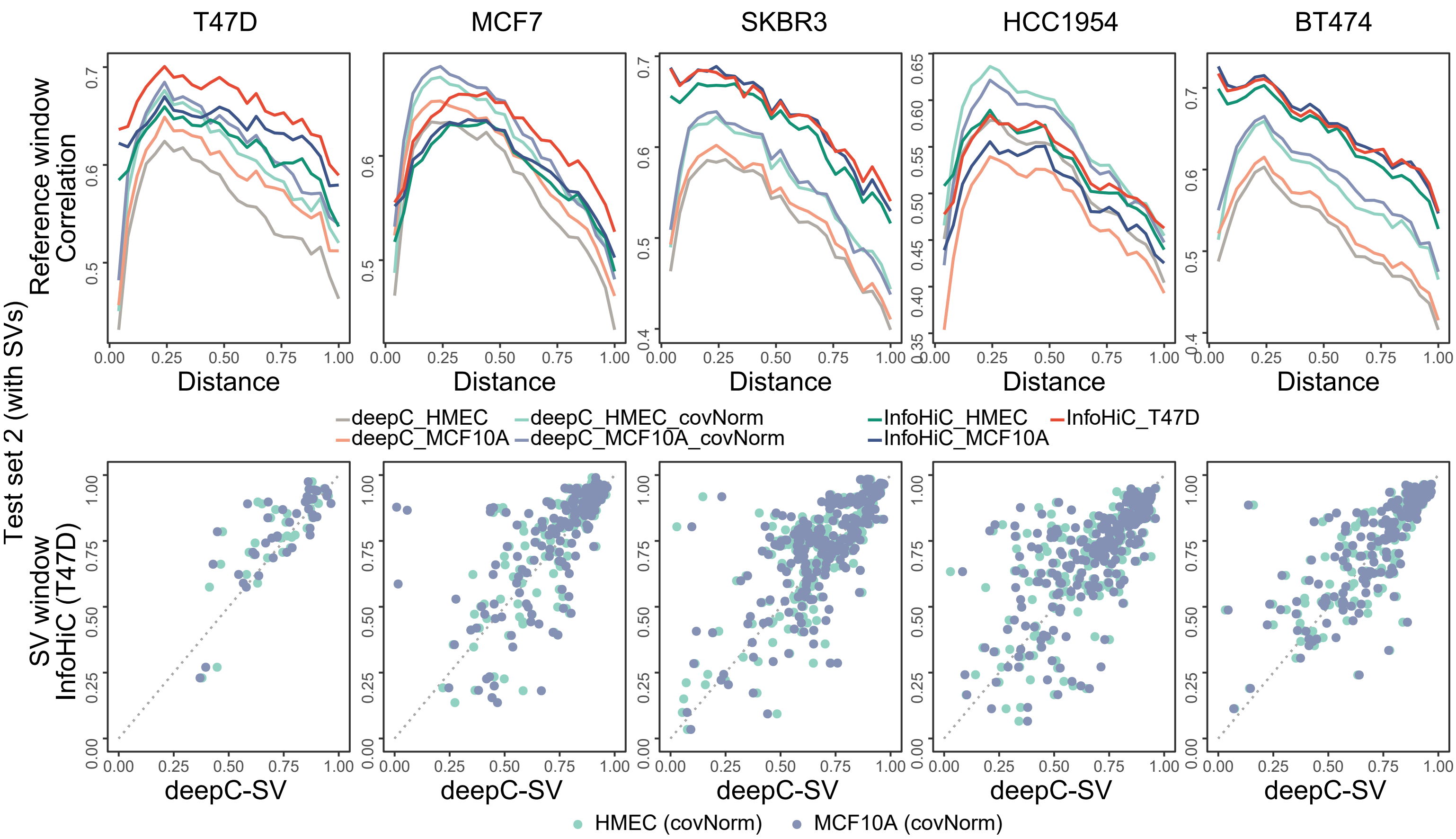


**Supplementary Fig. 1: Performance comparison between InfoHiC and deepC for the training usage of HMEC and MCF10A.** InfoHiC and deepC are compared for the training usage of the T47D, HMEC, and MCF10A cell line using the test set 2. The implicit normalization (covNorm) for deepC was applied to the HMEC and MCF10A Hi-C data. Distance-stratified correlations (line) for reference windows are shown (top) according to the mega-base scale distance range (0-2 Mb), and Pearson correlation (dot) per each SV window (bottom) is compared between the best models, deepC-SV trained by implicitly-normalized HMEC or MCF10A Hi-C data (bottom) and InfoHiC trained by the raw T47D Hi-C data (left).

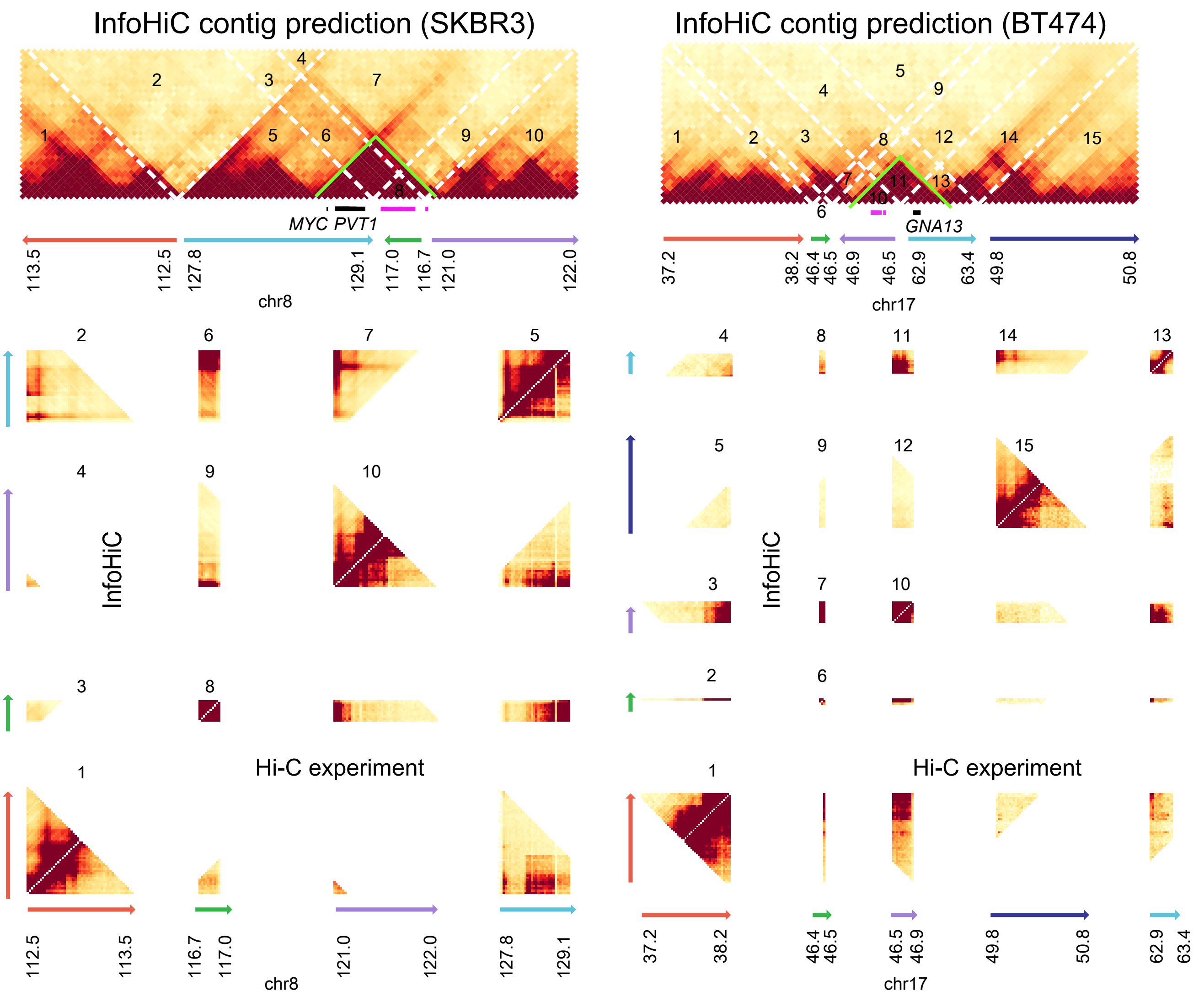


**Supplementary Fig. 2: InfoHiC validation scheme for examples of neo-TADs and SE hijacking events.** The contig Hi-C matrix has SV-derived contacts and each SV window is numbered. Dotted white lines represent SV breakpoints. Reference coordinates are shown below the contig Hi-C matrix at the megabase scale with the gene (black) and SE (crimpson) annotation. The total Hi-C matrix on the reference coordinate is shown at the bottom. The upper diagonal matrix is InfoHiC prediction and the lower diagonal matrix is the Hi-C experiment of the corresponding cell ilne. SV windows numbered from the contig Hi-C matrix are compared with Hi-C observation in the reference coordinate.

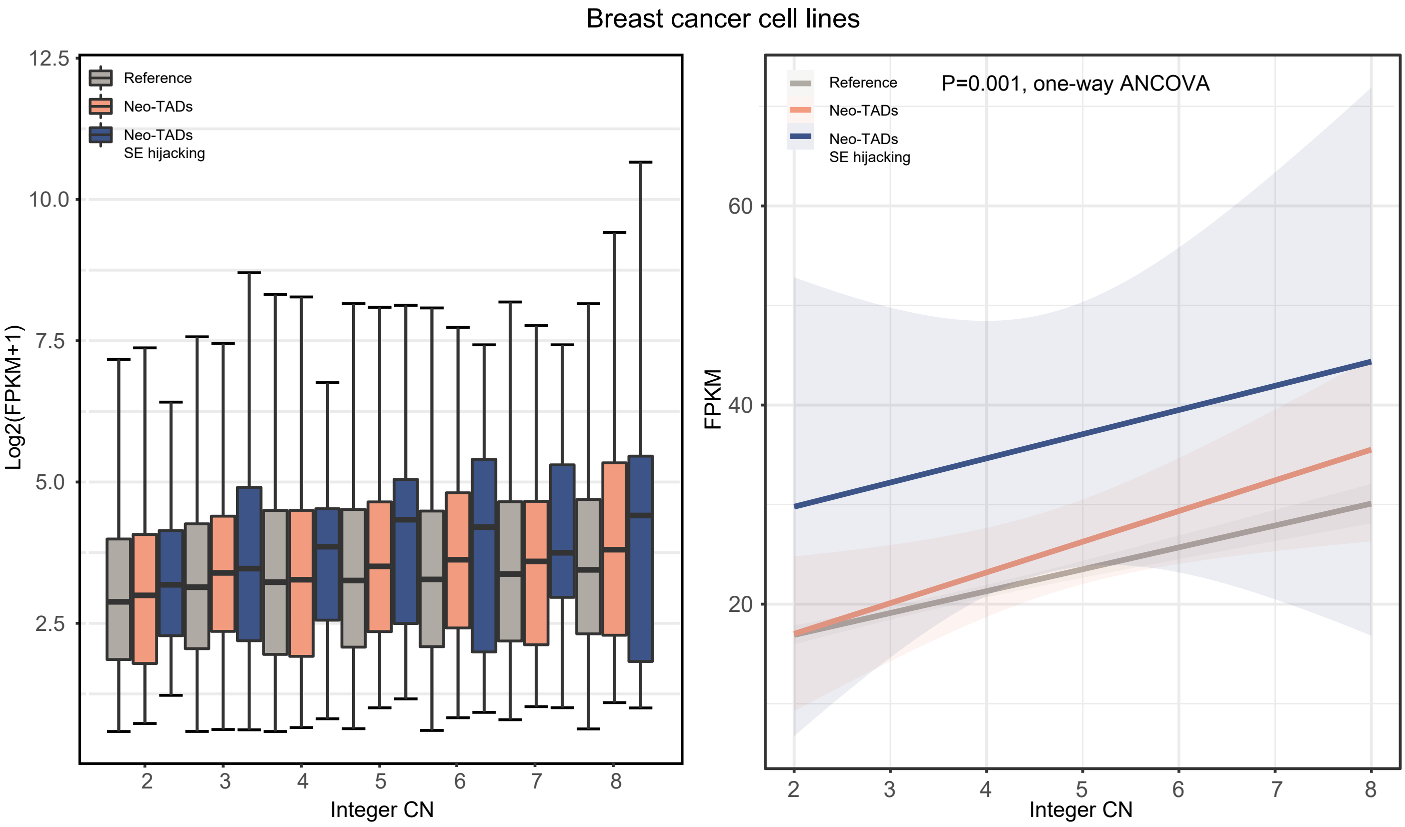


**Supplementary Fig. 3: Gene expression according to integer CNs in breast cancer cell lines.** Boxplots of RNA-seq Log2(FPKM+1) values depending on neo-TAD classes are shown according to integer CNs (left). The boxplot centre lines are medians, box limits are upper and lower quantiles, and wihskers are 1.5x interquantile ranges. The regression line between integer CNs and FPKM values is shown with the confidence interval for each neo-TAD class, and the P value was calculated by the one-way ANCOVA.

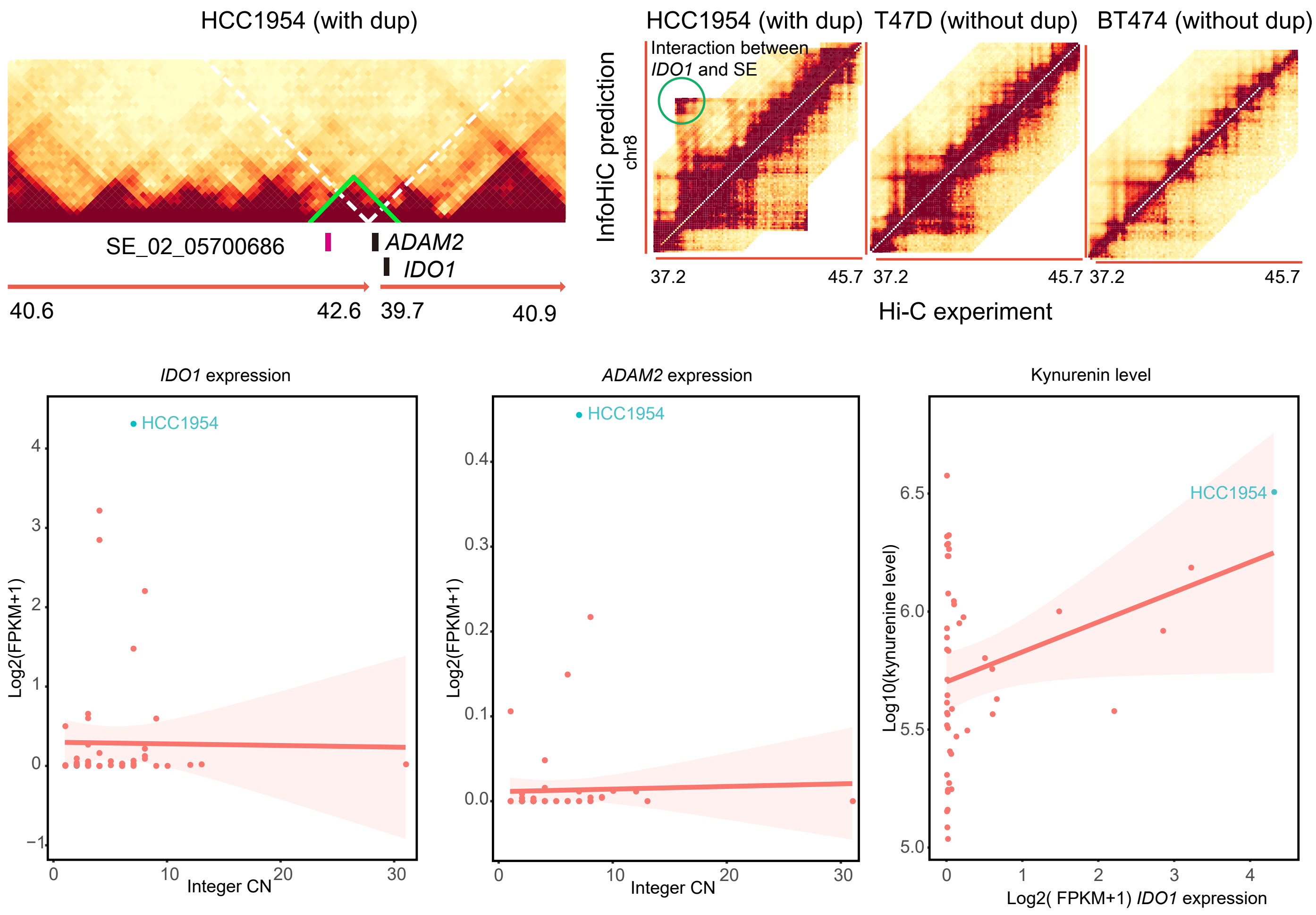


**Supplementary Fig. 4: The SE hijacking event of the *IDO1* gene in the HCC1954 cell line.**. The tandem duplication forms a neo-TAD (green) and results in the SE hijacking (crimpson) of the *IDO1* and *ADAM2* gene. In the reference coordinate, the InfoHiC prediction of the SE hijacking event was validated by the HCC1954 Hi-C experiment, and it was a novel interaction that is not observed in other cell lines without the duplication (T47D and BT474). Log2(FPKM+1) values of the *IDO1* and *ADAM2* gene are plotted versus integer CNs, and log10 values of the kynerenine level are plotted versus Log2(FPKM+1) value of the *IDO1* gene. Regression lines are shown with confidence intervals (red), and values from the HCC1954 cell line are shown in cyan.

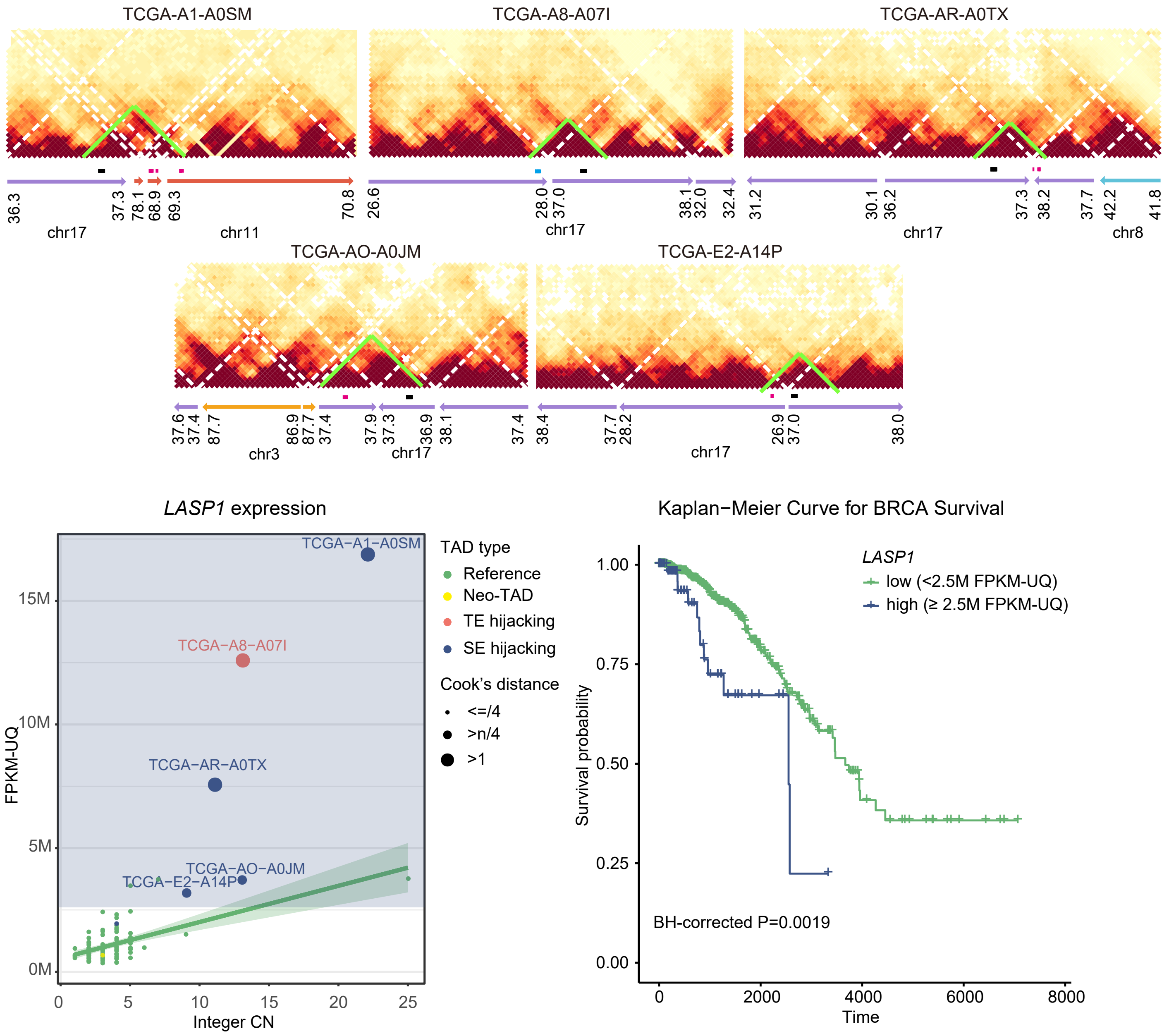


**Supplementary Fig. 5: InfoHiC prediction of contig Hi-C matrices of the *LASP1* region in BRCA patients.**. Neo-TADs are shown in green with hijacked SEs (crimson), hijacked TEs (blue), and the *LASP1* gene (black). Gene expression of *LASP1* of BRCA patients (dot) with WGS data (n=90). A linear regression line between FPKM-UQ values and integer CNs is shown with a confidence interval range (green). Neo-TADs are shown in different colors according to TAD types and in a different circle size according to the Cook's distance. The cutoff region of the FPKM-UQ value for overexpression is colored in the backgound (blue). Kaplan-Meier curve for BRCA survival with RNA-seq data (n=1098). The P value was calculated by the log-rank test, and adjusted by the BH procedure across neo-TAD overexpression genes.

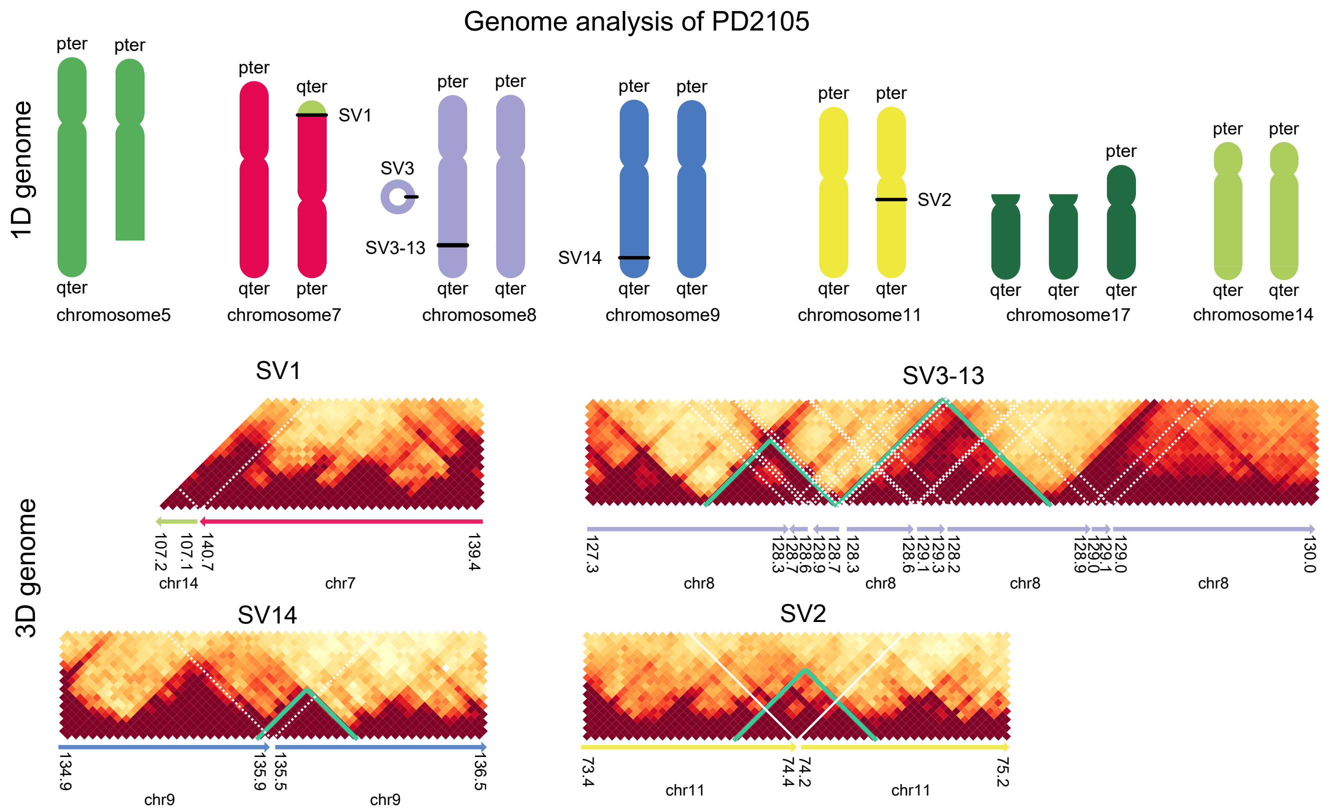


SV annotation

| Index | Chr1 | Pos1 | Chr2 | Pos2 | Direction | Type | Coding | Non-coding |
|---|---|---|---|---|---|---|---|---|
| SV1 | 7 | 140725137 | 14 | 107060339 | 3to5 | Tranlocation | | |
| SV2 | 11 | 74212451 | 11 | 74375379 | 5to3 | Duplication | Amplification of POLD3,RANP3 | |
| SV3 | 8 | 128231851 | 8 | 129281861 | 5to3 | Complex SV cluster (HSR/DM) | Amplification of MYC, POU5F1B | SE hijacking of MYC, POU5F1B |
| SV4 | 8 | 128883051 | 8 | 129134415 | 3to5 | | | |
| SV5 | 8 | 128929953 | 8 | 129040523 | 3to5 | | | |
| SV6 | 8 | 128251803 | 8 | 128656797 | 3to3 | | | |
| SV7 | 8 | 128251846 | 8 | 128932032 | 5to5 | | | |
| SV8 | 8 | 128579137 | 8 | 129012931 | 5to5 | | | |
| SV9 | 8 | 128730194 | 8 | 128930061 | 5to5 | | | |
| SV10 | 8 | 128864348 | 8 | 129040511 | 3to3 | | | |
| SV11 | 8 | 128932004 | 8 | 128965032 | 3to3 | | | |
| SV12 | 8 | 128579132 | 8 | 128867900 | 3to5 | | | |
| SV13 | 8 | 128965035 | 8 | 129134305 | 5to3 | | | |
| SV14 | 9 | 135465324 | 9 | 135889405 | 5to3 | Duplication | Duplication of GTF3C4,DDX31,AK8, TSC1 | SE hijacking of GFI1B |

**Supplementary Fig. 6: 1D genome and 3D genome analysis of the PD2105 patient.** SVs (SV1-14) are annotated in karyotypes (1D genome) of the PD2105 patient. InfoHiC predicted the 3D genome from the 1D genome and found neo-TADs (green) resulted from SVs that are represented by dotted white lines. Reference coordinates are shown at the megabase scale under the contig Hi-C matrices. SVs are classifed into simple and complex types, and their coding and non-coding effects are summarized in the SV annotation table.

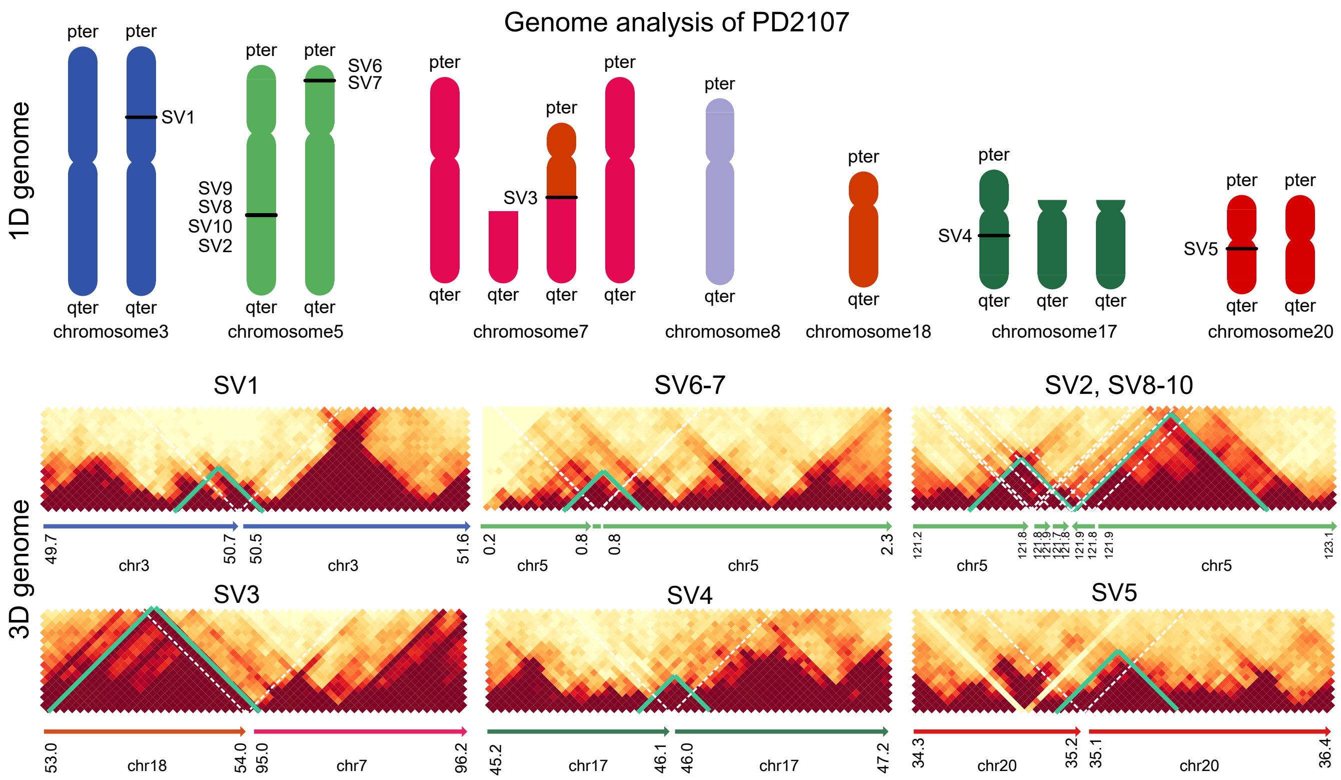


SV annotation

| Index | Chr1 | Pos1 | Chr2 | Pos2 | Direction | Type | Coding | Non-coding |
|---|---|---|---|---|---|---|---|---|
| SV1 | 3 | 50480341 | 3 | 50696608 | 5to3 | Duplication | Amplification of C3orf18,HEMK1, CISH, and MAPKAPK3 | |
| SV3 | 7 | 95044286 | 18 | 54010505 | 5to3 | Unbalanced translocation | | |
| SV4 | 17 | 46005379 | 17 | 46144955 | 5to3 | Duplication | Amplification of COPZ2,NFE2L1, PNPO, CDK5RAP3,PRR15L, MIR152, and NFE2L1-DT | |
| SV5 | 20 | 35080513 | 20 | 35190516 | 5to3 | Duplication | Amplification of DLGAP4,MYL9, and DLGAP4-AS1 | |
| SV6 | 5 | 701050 | 5 | 792312 | 3to5 | Insertion | | |
| SV7 | 5 | 696893 | 5 | 788253 | 5to3 | | | |
| SV2 | 5 | 121670510 | 5 | 121857025 | 5to3 | Complex SV cluster | Amplification of SNCAIP | Super-enhancer hijacking of PRDM6 |
| SV8 | 5 | 121755843 | 5 | 121858045 | 5to5 | | | |
| SV9 | 5 | 121755771 | 5 | 121858640 | 3to3 | | | |
| SV10 | 5 | 121755150 | 5 | 121774504 | 5to3 | | | |

**Supplementary Fig. 7: 1D genome and 3D genome analysis of the PD2107 patient.** SVs (SV1-10) are annotated in karyotypes (1D genome) of the PD2107 patient. InfoHiC predicted the 3D genome from the 1D genome and found neo-TADs (green) resulted from SVs that are represented by dotted white lines. Reference coordinates are shown at the megabase scale under the contig Hi-C matrices. SVs are classifed into simple and complex types, and their coding and non-coding effects are summarized in the SV annotation table.

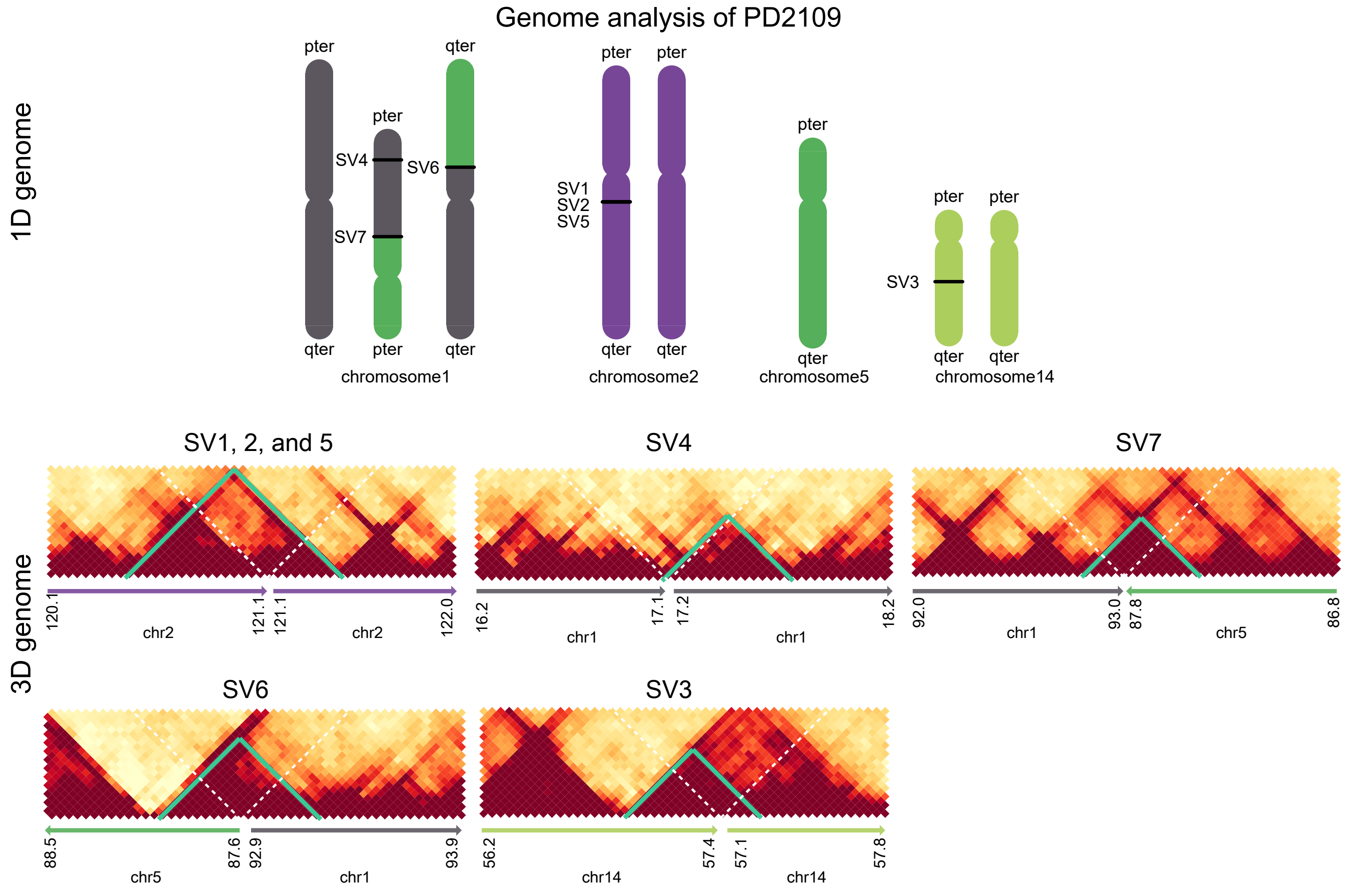


| Index | Chr1 | Pos1 | Chr2 | Pos2 | Direction | Type | Coding | Non-coding |
|---|---|---|---|---|---|---|---|---|
| SV1 | 2 | 121100120 | 2 | 121102933 | 5to5 | Complex SV cluster | | |
| SV2 | 2 | 121100344 | 2 | 121103431 | 3to3 | | | |
| SV5 | 2 | 121102544 | 2 | 121102940 | 5to5 | | | |
| SV6 | 1 | 92901932 | 5 | 87599726 | 5to5 | Reciprocal translocation | | SE hijacking of EVI5,GFI1 |
| SV7 | 1 | 92960263 | 5 | 87791087 | 3to3 | Reciprocal translocation | | SE hijacking of RPAP2,GFI1,GLMN |
| SV3 | 14 | 57069671 | 14 | 57401469 | 5to3 | Duplication | Amplification of TMEM260,OTX2 | |
| SV4 | 1 | 17054648 | 1 | 17189545 | 3to5 | Deletion | Deletion of MST1L | |

**Supplementary Fig. 8: 1D genome and 3D genome analysis of the PD2109 patient.** SVs (SV1-7) are annotated in karyotypes (1D genome) of the PD2109 patient. InfoHiC predicted the 3D genome from the 1D genome and found neo-TADs (green) resulted from SVs that are represented by dotted white lines. Reference coordinates are shown at the megabase scale under the contig Hi-C matrices. SVs are classifed into simple and complex types, and their coding and non-coding effects are summarized in the SV annotation table.

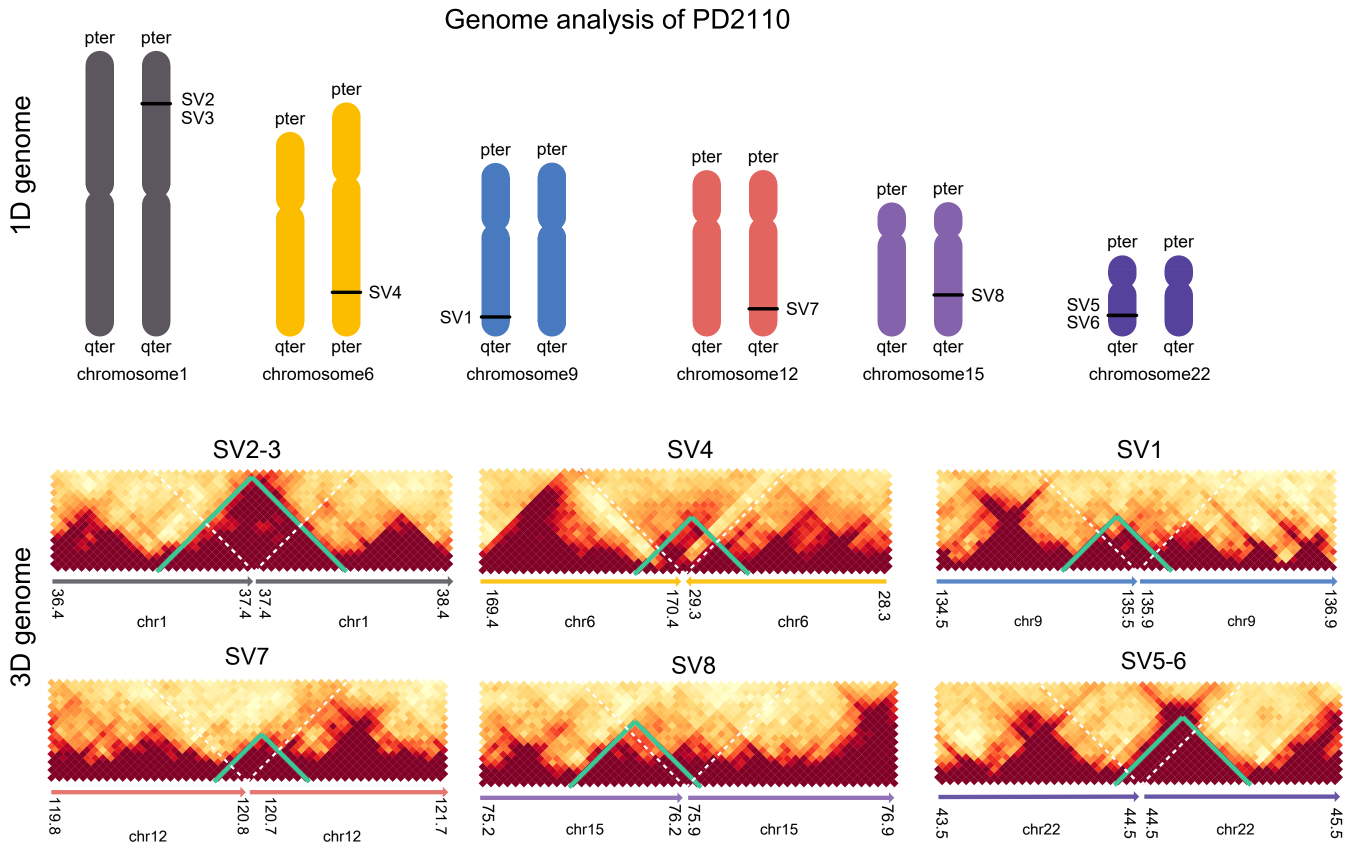


SV annotation

| Index | Chr1 | Pos1 | Chr2 | Pos2 | Direction | Type | Coding | Non-coding |
|---|---|---|---|---|---|---|---|---|
| SV1 | 9 | 135515011 | 9 | 135853084 | 3to5 | Deletion | Deletion of GTF3C4,DDX31,AK8, TSC1 | SE hijacking of SETX,TTF1,DDX31, BARHL1,GTF3C5, RALGDS,CEL, GFI1B |
| SV2 | 1 | 37423057 | 1 | 37425492 | 3to5 | Insertion | | |
| SV3 | 1 | 37423210 | 1 | 37425631 | 5to3 | Insertion | | |
| SV4 | 6 | 29290776 | 6 | 170356220 | 3to3 | Complex SV | | |
| SV5 | 22 | 44510365 | 22 | 44511823 | 3to5 | Insertion | | |
| SV6 | 22 | 44511438 | 22 | 44511863 | 5to3 | Insertion | | |
| SV7 | 12 | 120705648 | 12 | 120819844 | 5to3 | Duplication | Amplification of SIRT4,MSI1,PLA2G1B | |
| SV8 | 15 | 75899998 | 15 | 76189748 | 5to3 | Duplication | Amplification of UBE2Q2,SNUPN,CSPG4, SNX33,IMP3,ODF3L1 | |

**Supplementary Fig. 9: 1D genome and 3D genome analysis of the PD2110 patient.** SVs (SV1-8) are annotated in karyotypes (1D genome) of the PD2110 patient. InfoHiC predicted the 3D genome from the 1D genome and found neo-TADs (green) resulted from SVs that are represented by dotted white lines. Reference coordinates are shown at the megabase scale under the contig Hi-C matrices. SVs are classifed into simple and complex types, and their coding and non-coding effects are summarized in the SV annotation table.

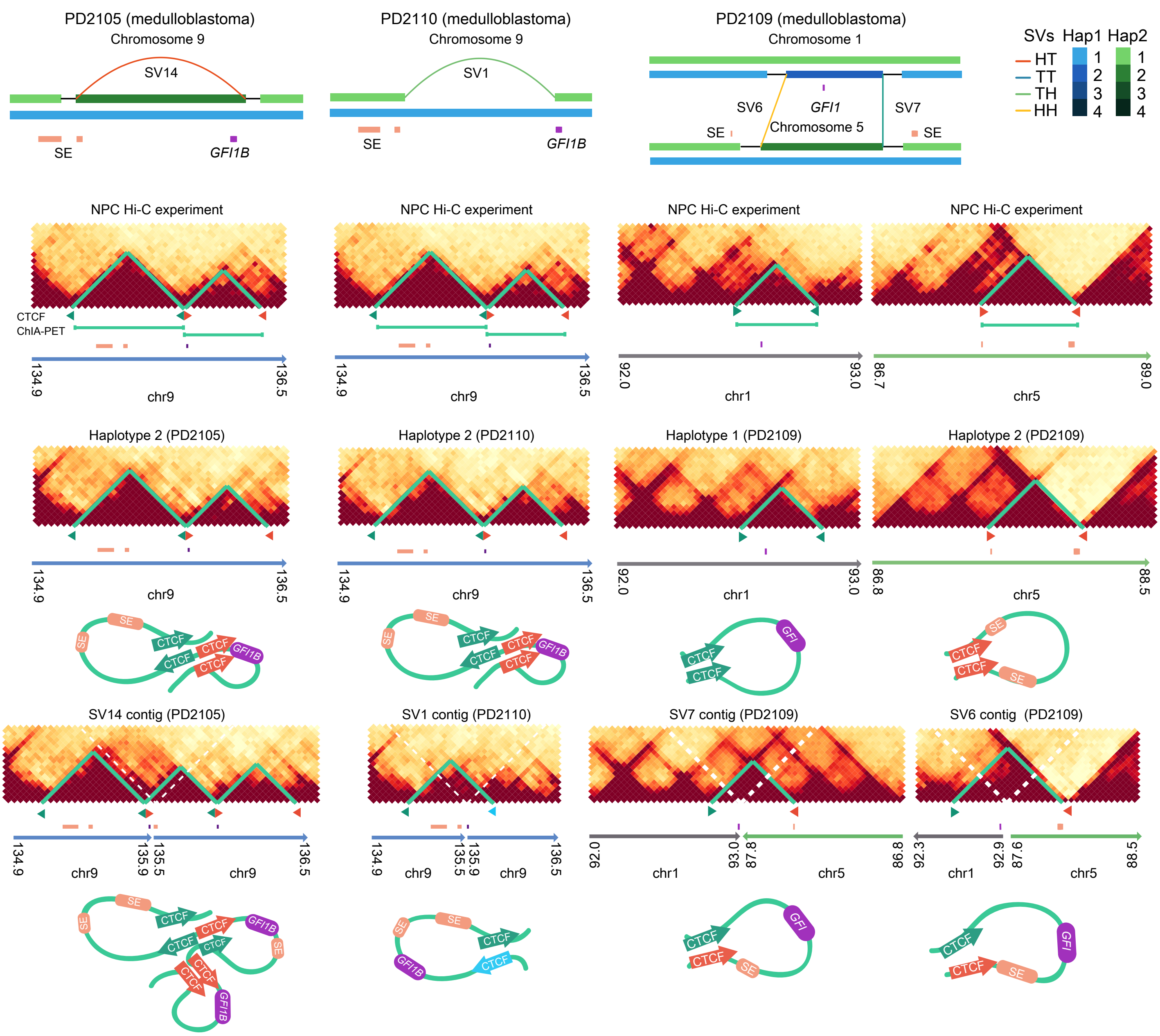


**Supplementary Fig. 10: SE hijacking events of *GFI1* gene families in patients with medulloblastoma.** Haplotype graphs of the patients with medulloblastoma and Hi-C experiments of the corresponding regions in neural progenitor cells (NPCs) are shown at the top. Hi-C matrices of contigs without or with SVs were predicted by InfoHiC. Reference TADs and neo-TADs are annotated (green) with CTCF motifs (arrowheads) near the TAD boundaries, and *GFI1B* and *GFI1* (purple) and SE (apricot) are annotated below. The pairwise interactions of the CTCF ChiA-PET data are indicated by green lines. 3D genome folding structures were modeled based on TAD boundaries and CTCF orientations.
